# Spatio-Temporal Dynamics of Oscillatory Brain Activity during the Observation of Actions and Interactions between Point-light Agents

**DOI:** 10.1101/2022.08.16.504090

**Authors:** Elisabeth V. C. Friedrich, Imme C. Zillekens, Anna Lena Biel, Dariusz O’Leary, Johannes Singer, Eva Victoria Seegenschmiedt, Paul Sauseng, Leonhard Schilbach

## Abstract

Predicting actions from nonverbal cues and using them to optimize one’s response behavior (i.e., interpersonal predictive coding) is essential in everyday social interactions. We aimed to investigate the neural correlates of different cognitive processes evolving over time during interpersonal predictive coding. Thirty-nine participants watched two agents depicted by moving point-light stimuli while an electroencephalogram (EEG) was recorded. One well-recognizable agent performed either a ‘communicative’ or an ‘individual’ action. The second agent either was blended into a cluster of noise dots (i.e., present), or was entirely replaced by noise dots (i.e., absent), which participants had to differentiate. EEG amplitude and coherence analyses for theta, alpha and beta frequency bands revealed a dynamic pattern unfolding over time: Watching communicative actions was associated with enhanced coupling within medial anterior regions involved in social and mentalizing processes and with dorsolateral prefrontal activation indicating a higher deployment of cognitive resources. Trying to detect the agent in the cluster of noise dots without having seen communicative cues was related to enhanced coupling in posterior regions for social perception and visual processing. Observing an expected outcome was modulated by motor system activation. Finally, when the agent was detected correctly, activation in posterior areas for visual processing of socially-relevant features was increased. Taken together, our results demonstrate that it is crucial to consider the temporal dynamics of social interactions and of their neural correlates to better understand interpersonal predictive coding. This could lead to optimized treatment approaches for individuals with problems in social interactions.

## INTRODUCTION

Social interactions build the foundation of our daily lives. Deficits in interpersonal relations have a negative impact on us and are associated with psychiatric disorders (Leong et al., 2021). It is as important to investigate, as it is difficult to research. Various brain areas and networks have been identified to be involved in social perception and interaction (Lahnakoski et al., 2012; Deen et al., 2015; Sliwa and Freiwald, 2017; Feng et al., 2021). Approaches to study social behavior range from passive observation of social interactions to interactive participation in social processes (Schilbach et al., 2013; Kingsbury and Hong, 2020; Misaki et al., 2021; Krol and Jellema, 2022). The latter form is closer to real-world social encounters, whereas the former has the advantage of investigating basic mechanisms underlying complex social interactions easier. Observing two persons interacting requires a continuous analysis of their bodies to be able to understand their non-verbal communication (Georgescu et al., 2014; Quadflieg and Koldewyn, 2017). Non-social (i.e., individual action) and social behaviors (i.e., communicative action) elicit different patterns in the observer’s brain (Isik et al., 2017; Wurm et al., 2017; Walbrin and Koldewyn, 2019). One way to investigate the perception of (non-) social behavior is to use point-light displays of agents, i.e., humans represented only by moving dots, which are attached to their major joints (Pavlova, 2012). Several studies showed that participants perceive point-light agents as biological motion (Neri et al., 1998; Grossman et al., 2000; Giese and Poggio, 2003; Peuskens et al., 2005). However, our brains do not only process bottom-up sensory information, but our perception and actions are influenced by our prior experience and might be modulated by social factors (Cracco et al., 2022). Our brain forms expectations and predicts mental states (Thornton et al., 2019), intentions (Kilner et al., 2007) and actions (Neri et al., 2006) of others. Specifically, we interpret nonverbal cues from one person to infer upon the presence and consequent action of another person to whom the behavior is directed. This phenomenon has been described as *interpersonal predictive coding* (Manera, Becchio, *et al*., 2011; Manera, del Giudice, *et al*., 2011; von der Lühe *et al*., 2016). According to the idea of interpersonal predictive coding, the brain generates hypotheses about subsequent actions of a potential interaction partner when seeing one person’s communicative gesture. When we see someone wave hello, we expect to see a second person, to whom this action is addressed. These interpersonal predictions are then compared to sensory input to make out whether, indeed, a second person is present or not. This can be especially important in noisy environments, e.g. to facilitate the detection of a point-light agent blended into a cluster of noise dots (Manera, Becchio, *et al*., 2011).

In this study, we aimed to disentangle the different processes involved in interpersonal predictive coding by analyzing the observer’s oscillatory brain activity in different time segments during the presentation of point-light agents’ actions and interactions.

Zillekens and colleagues (2019) used point-light agents to investigate interpersonal predictive coding in a functional magnetic resonance imaging (fMRI) study. They showed neuro-typical study participants a well recognizable point-light agent (agent A) and either a second point-light agent blended into randomly moving noise dots (agent B) or no second agent at all, but only noise dots. First, participants watched agent A performing either an individual or a communicative gesture. Second, they focused on agent B’s response action blended into a cluster of moving noise dots, or on noise dots that replaced agent B in 50% of the trials. Then, the participants had to indicate with a button press whether agent B had been present or absent in the cluster of moving dots. Behavioral results were in line with previous studies showing that communicative gestures, in contrast to individual actions, facilitated the perception of a second agent (Manera, del Giudice, *et al*., 2011) and increased its detection probability (Manera, Becchio, *et al*., 2011). On the neural level, the amygdala was found to be functionally coupled to the medial prefrontal cortex for communicative actions, a key region of the so-called mentalizing system of the brain, whose role for social cognition has repeatedly been demonstrated (Redcay and Schilbach, 2019). In the context of individual actions, the amygdala conversely showed increased connectivity to fronto-parietal areas, previously implicated as part of the attention system of the brain (Zillekens *et al*., 2019).

Given its high spatial resolution, fMRI allows to localize specific cortical and subcortical brain regions contributing to an effect. However, a temporal differentiation of sub-processes from observing nonverbal cues to the decision, which button to press is not possible. Being able to analyze social interactions in distinct time segments is crucial, as different cognitive processes might be involved at different time points. Moreover, splitting the task into time intervals enables us to investigate the initial processing of social gestures without confounding consequent factors such as matching or mismatching of predictions, the presence or absence of a second agent or trial outcome (Wurm and Caramazza, 2019). It also allows us to tie the time segments to specific events in the trial such as the response and, thus, our analyses of brain activity are not confounded by behavioral differences in reaction time for example.

In order to exploit these advantages, we therefore used electroencephalography (EEG) in combination with the established task to analyze the temporal dynamics of the cortical brain locations indicated by Zillekens and colleagues (2019) during interpersonal predictive coding (i.e., *a priori* based regions of interests (ROIs)). Additionally, we ran a data-driven whole brain analysis in order to see if further brain areas show specific activation for certain cognitive processes at a given time point. As Zillekens et al. (2019) also found differences in the neurofunctional coupling between the communicative and individual condition using blood oxygenation level dependent (BOLD) brain measurements, we performed exploratory coherence analyses with the *a priori*-defined ROIs according to the study of Zillekens et al. (2019).

We hypothesized that different cognitive processes would be involved at different time points in this complex task of social interaction as used by Zillekens and colleagues (2019). To investigate our claim, we split the task into three different time segments indicative of different cognitive processes involved in the task:

In the time segment (1) before the onset of agent B’s response action, participants focused on the well-recognizable agent A and viewed either a communicative or an individual gesture. In contrast to an individual gesture, a communicative action allowed the participants to form an immediate expectation for how agent B (if it was present) would act in response. Other studies have shown that the expectation of an upcoming action from someone else in a predictable context leads to an activation of one’s own sensorimotor system cortex, which has been interpreted as potentially reflecting mirror neuron activity (Kilner *et al*., 2004; Maranesi *et al*., 2014; Krol *et al*., 2020). The mentalizing system on the other hand has also been proposed to be crucial to infer intentions from observed actions (Van Overwalle, 2009; Van Overwalle and Baetens, 2009; Hamilton and Marsh, 2013; Spunt and Lieberman, 2013; Catmur, 2015; Begliomini *et al*., 2017). Socially intended movements as well as imagined social interactions have been shown to lead to an activation in regions of both mirror neuron and mentalizing systems (Becchio *et al*., 2012; Trapp *et al*., 2014). Especially the posterior superior temporal sulcus – part of both systems-was suggested to be essential for the perception of social interactions, which provides the basis for social action understanding and interpretation (Isik *et al*., 2017; Walbrin *et al*., 2018). Besides the social content, the generation of predictions based on the communicative gesture might need more cognitive resources than in the individual condition. These differences in executive control and cognitive load could also lead to differences in prefrontal activation.

In the time segment (2) after the onset of agent B’s response actions (or its replacement with noise dots), participants were able to validate their expectations and tried to detect a second agent in the cluster of moving dots. Communicative actions in contrast to individual ones performed by the first agent were thought to allow specific predictions of agent B’s response action and should, thus, facilitate the detection of it (Manera, Becchio, *et al*., 2011; von der Lühe *et al*., 2016; Zillekens *et al*., 2019). This could again lead to differences in the cognitive resources needed between conditions. Moreover, an expected outcome should lead to more motor system activation than a non-expected outcome (Braukmann *et al*., 2017).

In the time segment (3) before the participants’ response, the participants had to decide whether the agent is actually present or absent. Inferring human actions from point-lights was shown to activate the motor system (Ulloa and Pineda, 2007; Perry *et al*., 2010). The motor system might also be required to fill in the gaps between the point-light dots, so the single points are perceived as biological motion and thus a second agent rather than noise is seen (Saygin et al., 2004). Therefore, we expected pre- and post-central ROIs to be activated more in signal than in noise trials.

Behaviorally, we used two signal detection theory parameters to differentiate between the presence and absence of a second agent (i.e., the sensitivity) and to evaluate the tendency to choose presence or absence (i.e., the response criterion) (Stanislaw and Todorov, 1999). We expected that participants would perform better when point-light agents engage in communicative actions rather than in individual ones (expressed by enhanced sensitivity in communicative trials) and would be more likely to perceive a second agent after communicative than individual gestures (expressed by a decreased, i.e. less conservative, response criterion in communicative trials) (cf. Manera, Becchio, *et al*., 2011; Manera, del Giudice, *et al*., 2011; Zillekens *et al*., 2019). As these previous studies suggest that communicative gestures provide information to predict the other person’s behavior and thus facilitate the perception of another agent, reaction times should be faster in communicative than in individual trials. Detecting an actually present agent B will result in an immediate response and thus signal trials are expected to result in faster reaction times than noise trials (with agent B being absent).

## METHODS

### Participants

An *a priori* power analysis was performed using G*power (Faul *et al*., 2007) to determine the appropriate sample size. We used a power of 0.95 and an estimated effect size dz of 0.56 based on the behavioral study of Zillekens et al. (2019), which showed that participants achieved a higher performance in trials with communicative compared to individual actions using the same experimental design. The *a priori* power analysis suggested a sample size of 36 when using a one-tailed t-test for matched pairs and a significance level of 0.05. As artifacts are common in EEG studies, we recorded 15% more participants, resulting in a sample size of 41 volunteers. Using Mueller-Putz et al.’s (2008) formula as a basis, two participants were removed from all analyses, as their performance did not surpass chance level significantly. The final sample consisted of 39 participants (21 female, 18 male) between the ages of 20 and 50 years (M = 25.62, SD = 6.37), all right-handed with normal or corrected to normal vision, no diagnosis of neurological or psychiatric disorders, and no history of medication intake. Additionally, Autism Spectrum Quotient (AQ) values were indicative of a neuro-typical control group (M = 15.64, SD = 5.55) (Baron-Cohen *et al*., 2001). AQ values ranged between 6 and 28, which means that they were all below the clinical cut-off value of 32 proposed by Baron-Cohen and colleagues (2001).

Flyers and circular emails were used to recruit volunteers at the Ludwig-Maximilians-Universität München. A monetary compensation of 10 € per hour was given to all participants for their time and effort. The study was approved by the ethics committee of the Faculty of Psychology and Education Sciences at Ludwig-Maximilians-Universität München and is in accordance with the guidelines of the Declaration of Helsinki. Some data from a sub-sample of the recorded participants has been published in relation to a research question not addressed in this manuscript (Friedrich et al., 2022).

### Experimental Design

The experimental design was controlled using the Psychophysics Toolbox (Version 3.0.14; Brainard 1997; Kleiner, Brainard, and Pelli 2007; Pelli 1997) in Matlab R2016a (The MathWorks, Inc., Natick, Massachusetts, United States) and was based on Zillekens et al.’s (2019) study.

Participants were facing a computer monitor (refresh rate = 60 Hz, resolution of 1280 × 1024) displaying black dots moving on a grey background (Figure 1). One half of the display contained moving dots that represented a well recognizable point-walker (agent A), executing either one of three communicative or one of three individual actions (communicative condition: asking to squat down, asking to look at the ceiling, asking to sit down; individual condition: turning around, sneezing, drinking). In 50 % of the trials, the other half of the display contained a second point-walker (agent B) within a cluster of spatially and temporally scrambled moving dots, which responded to agent A’s communicative action (i.e., signal trials: agent B present and squatting down, looking at the ceiling or sitting down). In the remaining 50 % of trials, agent B was substituted by randomly moving noise dots (i.e., noise trials: agent B absent). In order to make the visual input comparable between conditions, signal and noise trails always contained the same number of dots, either constituting an agent B or moving in a random fashion. Agent A’s actions therefore determined if a trial belonged to the communicative or individual condition, whilst agent B’s presence or absence defined a trial as a signal or noise trial (Figure 1A).

**Figure 1.**
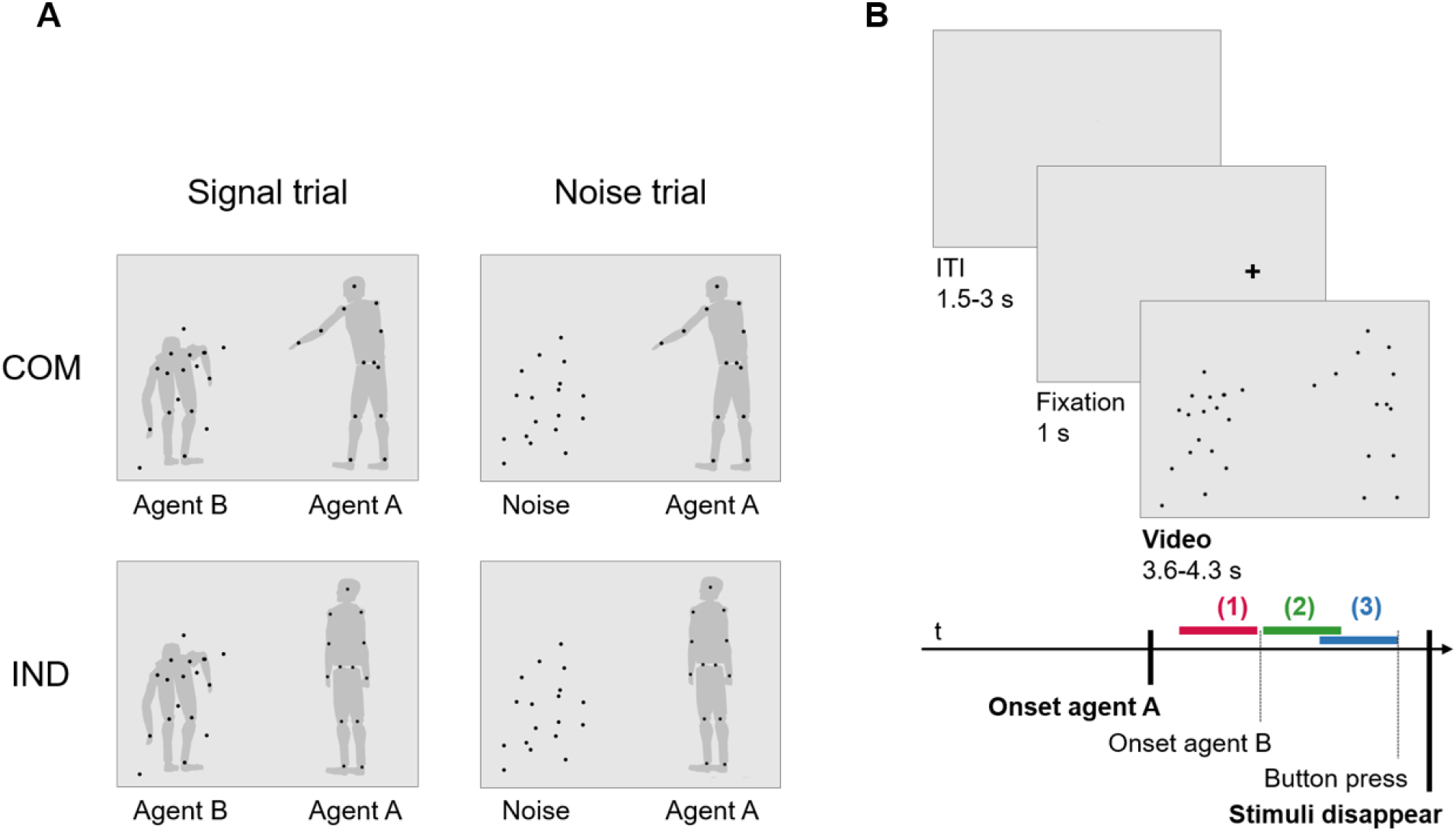
Experimental design (adapted from Zillekens et al. (2019) and Friedrich et al. (2022)). (A) Experimental conditions. Participants observed a screen with two point-light agents. Agent A either performed a communicative (COM, see top row) or an individual (IND, see bottom row) gesture. The responding agent B was either present and blended into a cluster of noise dots (i.e., signal trial, see left column) or was absent and entirely replaced by noise dots (i.e., noise trials, see right column). As the agents simply consisted of dots, they were only recognizable when performing a biological movement. As the movement of the dots could not be displayed in the figure, here, we have added grey silhouettes for display purposes only. Note that these silhouettes were not displayed in the experimental task for the participants to see (compare to Figure 1B in which the agents are illustrated as the participants saw them). (B) Structure of a trial. A trial consisted of a jittered inter-trial-interval between 1.5-3 s and a 1-s fixation cross indicating at which position agent A will appear next. Then the video with the two point-light agents started (i.e., ‘Onset of agent A’ written in bold). Although both agents were present right from the video’s onset, agent B’s response action was delayed. During the delay, agent B (and the added noise dots) was depicted by moving dots as though as it was swaying back and forth and ‘watching’ agent A. The exact onset of agent B’s goal-directed response action and the duration of the video varied between trials and was dependent on the actions performed. Participants were asked to first focus on agent A. Then they should observe the cluster of dots and decide by button press whether agent B was present or absent. They had to make their choice before the video with the two agents ended and the stimuli on the screen disappeared (i.e., ‘Stimuli disappear’ written in bold). We divided the video in three time segments for analyses. Time segment (1) before agent B’s onset: Participants passively observed agent A performing either a communicative or individual action (red color). Time segment (2) after agent B’s onset: Participants watched the responding agent B, which was blended into a cluster of noise dots or was entirely replaced by noise dots (green color). Time segment (3) before the response: Participants decided whether agent B was present or absent in the cluster of noise dots and indicated their choice with a button press (blue color).

Trials began with the presentation of a blank screen for a jittered inter-trial-interval between 1.5 – 3 s, followed by a fixation cross for 1 s indicating if agent A would appear on the left or right half of the display (Figure 1B). Following this, the video depicting agent A simultaneously with the cluster of dots on the opposing side of the display (with or without the presence of agent B) was shown. Although the dots constituting agent B were visible from the beginning, the onset of agent B’s response action was delayed. During the delay, agent B (and the added noise dots) was already represented by moving dots as though it was swaying back and forth and ‘watching’ agent A’s actions. Agent B’s response action was defined by a goal/spatially-directed dot movement exceeding a velocity of 10 angles/s for a minimal duration of 100ms (Ansuini *et al*., 2015). The delay between the start of agent A’s action and the start of agent B’s action, ranged between 1267 - 1567 ms and was dependent on the specific movement. The whole video had a duration ranging between 3600 - 4300 ms based on the particular action shown.

Participants were instructed to watch agent A first and only then attend to the cluster of dots. Following this, they were asked to press a button as swiftly as possible to indicate if agent B was present (signal) or absent (noise). Participants had to respond while the video was still on-going. Once the dots vanished, responses were treated as invalid. Using a standard German (QWERTZ) keyboard, response button assignment for yes and no (V- or N-key) was counterbalanced across participants. Furthermore, which point-walker appeared on the left or right side of the display was counterbalanced.

Analogous to the approach in Zillekens et al. (2019), the agents’ actions were selected from the Communicative Interaction Database (Manera *et al*., 2010). The actions were reliably identified as communicative or individual in their study (Manera *et al*., 2010). We chose the communicative actions ‘asking to squat down’, ‘asking to look at the ceiling’ and ‘asking to sit down’ because they were used in several studies relevant for our hypotheses (Manera, Becchio, *et al*., 2011; Manera, del Giudice, *et al*., 2011; Zillekens *et al*., 2019). Moreover, they were confirmed to be reliably detected as communicative actions and produced a difference in sensitivity between communicative and individual gestures (Manera, Becchio, *et al*., 2011). On the screen, the agents always faced each other and kept a comparable distance from the center. The number of dots, the movement velocity as well as the onset an duration of the movements were kept stable between the communicative and individual conditions (Manera *et al*., 2010). In the communicative condition, agent B’s response action matched the communicative action of agent A in timing, position and kinematics (Manera, del Giudice, *et al*., 2011). In the individual condition, agent A’s communicative movement was replaced by an individual action with an identical temporal onset and duration.

The dots constituting agent B were presented using a limited lifetime technique: Thirteen possible dot locations defined agent B’s body. However, at any given time only 6 of these possible locations comprised signal dots. Every 200 ms, a dot would disappear and reappear at another location, with the timing of dot appearance being desynchronized. This limited lifetime technique was implemented to prevent participants from depending on the simultaneous transition of dots representing agent B’s body. For more details, please see Manera, del Giudice, *et al*. (2011) and Zillekens *et al*. (2019).

### Procedure and EEG Data Acquisition

All participants provided written informed consent, their age, gender, and completed the short form of the Edinburgh Handedness Inventory (Veale, 2014) in addition to the online version of the AQ (Autism Research Center; Baron-Cohen et al. 2001) prior to the EEG experiment. Moreover, the absence of neurological or psychiatric disorders, brain injuries, medication intake and pregnancy were all confirmed in a short telephone interview.

To record EEG during the experiment, 61 scalp electrodes (Ag/AgCl ring electrodes; Easycap ®) were placed following the international 10-10 system. Electrodes were applied on the right and left outer canthi and below the left eye to measure the electrooculogram (EOG). Fpz was used for the ground electrode and the reference was positioned on the nose tip. A sampling rate of 1000 Hz and a 64-channel amplifier system (BrainAmp, Brain Products ®) were used for signal acquisition, while impedances were maintained below 10 kΩ.

To ensure participants could achieve an average performance around 70% correct responses, an established procedure utilizing a pretest of 108 trials was employed to individually adapt the difficulty level (i.e., the number of noise dots added to agent B); The screen showed a cluster of dots and after its disappearance, participants had to press a button to indicate whether agent B was present or absent within the cluster of dots. In contrast to the main experiment, there was no agent A displayed in the pretest but only the cluster of dots (with or without agent B) with alternating difficulty (i.e., 5, 20 or 40 added noise dots). By fitting a cumulative Gaussian function to the performance of the participants the number of dots necessary for an accuracy of 70% was calculated. A minimum of five dots was used, regardless of if the calculated number of dots was lower (Manera, Becchio, *et al*., 2011; Manera, del Giudice, *et al*., 2011; Zillekens *et al*., 2019). The resulting number of noise dots for each individual participant was added to agent B in the main experiment to make its detection 70% accurate.

As the main experiment (see Figure 1) differed from the pretest, participants were given the opportunity to practice the experimental task during twelve practice trials. Following this, participants completed 288 experimental trials divided into four 9-min blocks with breaks included between blocks.

### Behavioral Data Analyses

Behavioral data analyses were completed in Matlab R2016a, whereas statistical analyses were performed in IBM® SPSS® Statistics version 24.0.0.0 and JASP version 0.9.2 and 0.16.3 (JASP Team, 2019).

We defined the onset of agent B’s response action at 1567 ms after the video onset for the action ‘sitting down’, 1500 ms for the action ‘squatting down’, and 1267 ms for the action ‘looking at ceiling’ (Figure 1B) (Ansuini *et al*., 2015).

Trials in which a response was given prior to the onset of agent B plus 200 ms or after agents A and B had already disappeared were removed from the analysis. Reaction times comprised the time window between the onset of agent B’s response action and the button press (Figure 1B).

We calculated the hit rate (proportion of true positive responses of all signal trials) and the false alarm (FA) rate (proportion of false positive responses of all noise trials), z-transformed the rates and computed the two signal detection theory parameters sensitivity d’ and response criterion c as follows (Macmillan and Creelman, 1990; Zillekens *et al*., 2019):

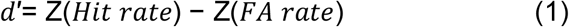

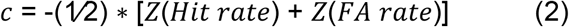

### EEG Data Analyses

BrainVision Analyzer 2.0 (Brain Products ®) was employed to preprocess the EEG data. The whole preprocessing was carried out on continuous EEG data. A preliminary visual inspection was conducted to remove large artifacts and interpolate bad channels. Out of the 61 EEG channels, 1 channel had to be interpolated for 3 participants, 2 channels for 2 participants and 3 and 5 channels each for one participant. Data was filtered between 0.1 and 120 Hz (48 db/oct), and a notch filter was applied at 50 Hz. Re-referencing of EOG channels was accomplished using a bipolar reference (i.e., right – left horizontal EOG channels, bottom vertical EOG channel – FP1), whereas a common average reference was utilized for the re-referencing of EEG channels. Blinks (vertical eye movements) and saccades (horizontal eye movements) were identified with an Independent Component Analysis for ocular correction on the bipolar EOG channels. Components that clearly represented artifacts were removed, on average 6.5 components (SD = 3.7) were removed per participant. Then, a further visual inspection was conducted to identify and remove any remaining artifacts. To accommodate the Standardized Low Resolution Electromagnetic Tomography (sLORETA) analyses data was re-sampled to 1024 Hz.

After the preprocessing, the continuous data was segmented: As the onset of agent B and the time point of participants’ responses varied between trials, data could not be averaged and analyzed over the entire time course. Instead, each trial was separated into three 1-s long time segments in relation to these two events (Figure 1B):

1. From 1-s before agent B’s onset to agent B’s onset (i.e., observation of agent A)
2. From agent B’s onset to 1-s after agent B’s onset (i.e., observation of agent B)
3. From 1-s before the response to the response (i.e., decision whether agent B is present or absent)

Solely trials in which a response was given 1 s after agent B had started its response action and ahead of the stimuli disappearing were included. An average of 25 trials (SD= 23.4) of the total 288 per participant had to be discarded because the response occurred before 1 s. An average of 37 trials (SD= 24.8) had to be discarded because no response or a response occurred after the agents had disappeared. All analyzed trials are free of motor responses, while simultaneously containing a valid response. As the mean reaction time over participants and all trials was 1601 ms, time segment (2) and (3) overlapped on average about 400 ms.

Segments without artifacts were exported for the following contrasts:

1. communicative (i.e., COM: agent A performs a communicative action) versus individual (i.e., IND: agent A performs an individual action) condition
2. non-expected (i.e., trials in which a communicative action from agent A is followed by noise or an individual action from agent A is followed by the presence of agent B) versus expected outcome (i.e., trials in which a communicative action from agent A is followed by the presence of agent B or an individual action from Agent A is followed by noise).
3. signal (i.e., agent B was present) versus noise (i.e., agent B was replaced by randomly moving noise dots) trials

#### Source-level Amplitude Analyses

The data was further analyzed using Matlab R2015b and sLORETA v20190617 (Pascual-Marqui, 2002; Pascual-Marqui, 2007). To avoid confounding factors from incorrect responses, we only analyzed trials with correct responses: Signal trials in our neural analyses, therefore, mean that agent B was present and was indeed correctly detected (i.e., hit). Noise trials mean that agent B was not present and was also not erroneously detected (i.e., correct rejection).

After discarding all trials with artifacts, invalid (i.e., response was too early or too late, see previous paragraph) and false responses, we only analyzed participants who still had ≥ 20 remaining trials for each condition. All 39 participants had ≥ 20 trials for the contrast communicative versus individual condition (on average 79 trials per condition) and for the contrast signal (on average 85) versus noise trials (on average 73 trials). For the contrast of non-expected versus expected outcome, 4 subjects had less than 20 trials for each condition, thus this contrast only included 35 participants. These participants had on average 36 trials for the communicative noise trial (COM-Noise), 43 for the individual signal trial (IND-Signal), 45 for the communicative signal trial (COM-Signal) and 38 for the individual noise trial (IND-Noise). Please refer to the Table S1A for more details about the number of trials (mean, standard deviation and range per condition).

The amplitude analysis comprised the computing of EEG crossspectra across all single trials, for each individual participant and condition, in the frequency range between 1-40 Hz, and the extraction of global field power. Subsequently, the scalp-level data was converted into voxel-based sLORETA data (i.e., source space).

sLORETA (i.e., Standardized Low Resolution Electromagnetic Tomography) is a method to transfer electrical activity measured by scalp electrodes into source space (Pascual-Marqui, 2002; Pascual-Marqui, 2007). It solves the inverse problem (i.e., inferring the brain sources from scalp electrode potentials) and localizes the sources of the brain activity. It was shown that an exact source reconstruction is possible using LORETA with 64 scalp electrodes (Akdeniz, 2016).

For the *a priori* ROI-based analyses, the data for each of the contrasts was extracted for the coordinates of the specific ROIs according to the study of Zillekens et al. (2019), which showed a significant difference in the BOLD signal within the three contrasts and were close enough to the cortical surface to be recorded with EEG (see Supplementary Tables S2-S4). For the data-driven whole brain analyses, this step was not necessary, as all voxel-based data entered the statistical analysis.

The statistical analyses for the *a priori* ROI-based and the data-driven whole brain data were performed over the (A) theta (4-7 Hz), (B) alpha (8-12 Hz), (C) lower beta (13-17 Hz), and (D) upper beta (18-25 Hz) frequency band for each of the three time segments: 1-s before and 1-s after agent B’s onset and 1-s before the response. To compute a two-tailed t-statistic for paired groups the sLORETA built-in voxelwise randomization test (5000 permutations) was employed. This method to correct for multiple comparisons relies on statistical non-parametric mapping to calculate corrected critical thresholds and p values.

#### Scalp-level Amplitude Analyses

The scalp-level amplitude analyses had the purpose to evaluate whether the significant source-level effects can be replicated on scalp-level. The scalp-level amplitude analyses were also performed in sLORETA and were based on the same trials, EEG crossspectra and extracted field power between 1-40 Hz that were calculated for the source-level amplitude analyses. Instead of transforming the data into source space, the scalp electrode was selected that fits best to the underlying significant source-level effect according to the study of Scrivener and Reader (2022).

The statistical analyses were made for the chosen electrode and the contrasts that became significant in a specific time segment and frequency band in the source-level. We computed one-tailed t-statistic for paired groups with the sLORETA built-in voxelwise randomization test (5000 permutations) to correct for multiple comparisons and calculate corrected critical thresholds and p-values.

#### Source-level Coherence Analyses

We performed exploratory coherence analyses only for the contrast communicative versus individual condition based on the *a priori*-defined ROIs according to the study of Zillekens et al. (2019) (Supplementary Table S5). These ROIs showed differences between the communicative and individual condition in their functional coupling to the amygdala. Here, we investigated differences between the communicative and individual condition in their functional coupling to each other (because it is not possible to capture the amygdala’s activity with EEG).

For the coherence analyses, the preprocessing, the number of participants and the analyses tools were the same as for the source-level amplitude analyses. The figures were created using Fieldtrip (Tzourio-Mazoyer et al., 2002; Oostenveld et al., 2011). We also used only correct trials and the same minimum number of trials. However, for the coherence analyses, the number of trials between conditions was kept equal for each participant (as unequal numbers of trials across conditions can influence coherence values). This was done by randomly excluding trials until the lowest number of trials in any condition was matched. In the end, participants had 76 trials per communicative and individual condition (see Table S1B for more details).

For each condition, EEG crossspectra were computed over the remaining single trials in the frequency range between 1-40 Hz, separately for each subject and condition. Lagged coherence values (eliminating any spurious zero-phase lag coherence due to volume conduction) were extracted for the defined ROIs in source space according to Supplementary Table S5. Coherence coefficients vary between 0 and 1. Two-tailed, paired t-statistics for theta, alpha, lower beta and upper beta frequency bands were computed for all possible ROI connections. Correction for multiple comparisons was applied using randomization tests the same way as done for the amplitude analysis.

## RESULTS

### Behavioral Results

As intended by the individual adjustment of difficulty, participants achieved a performance of 71.65% (SD = 8.66) correct responses at an average difficulty level of 13.05 (SD = 10.07; range = 5 -44) interfering noise dots.

Normal distribution was not violated for the sensitivity (d’), thus a one-tailed paired-sample t-test was calculated to test the hypothesis of higher sensitivity d’ in the communicative as compared to the individual condition. The increase in d’ in the communicative (M = 1.32, SD = 0.61) compared to the individual condition (M = 1.29, SD = 0.64) remained non-significant (t(38) = 0.38, p = .35; dz = 0.061; Figure 2A).

**Figure 2.**
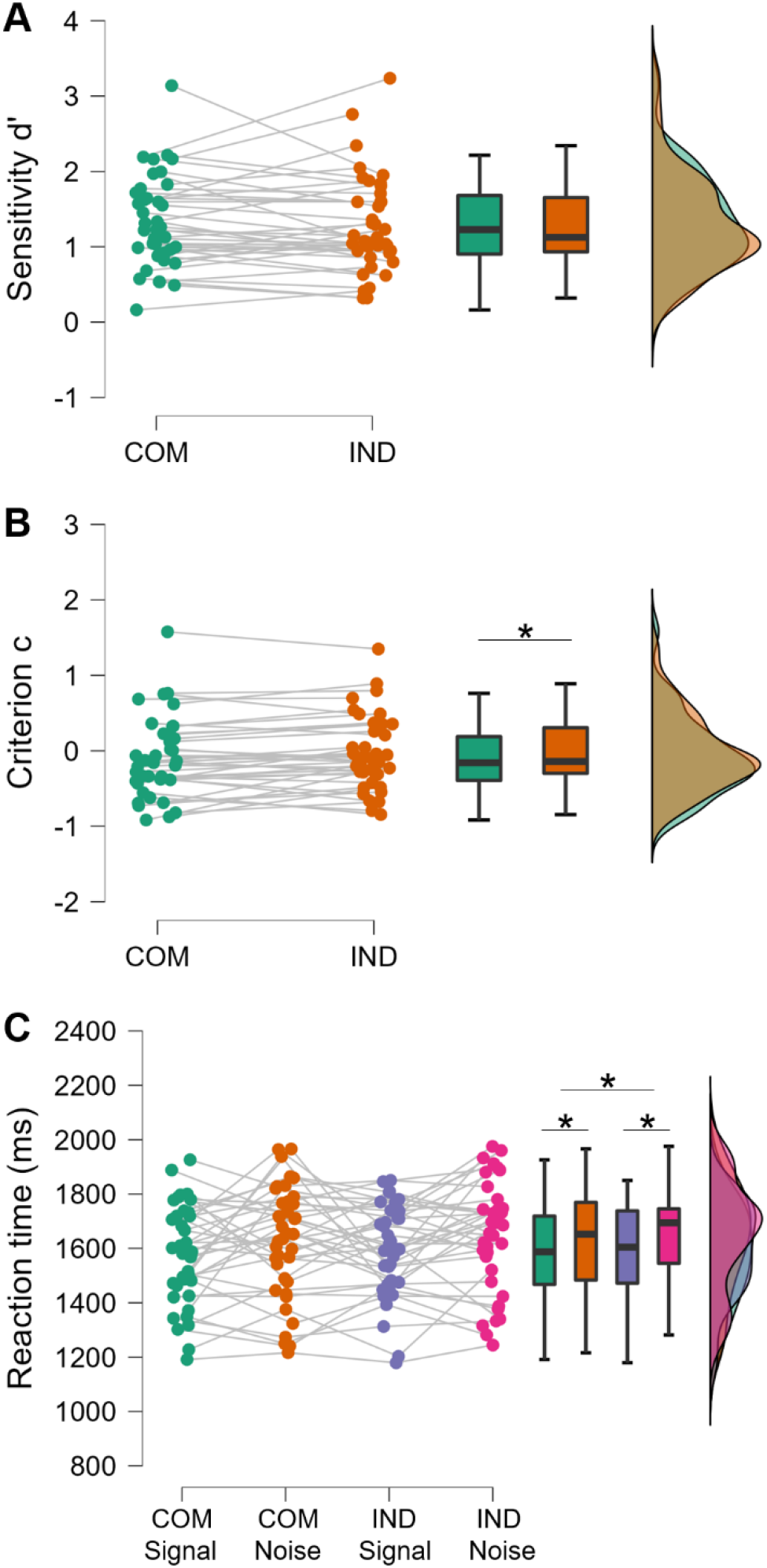
Behavioral results. (A) Mean sensitivity d’ values and (B) mean criterion c values in the communicative (COM) and individual (IND) conditions. (C) Mean reaction times in ms for signal and noise trials separately for the communicative and individual condition. The raincloud plots show first the individual data points, then box plots and last the distribution of the data. Statistically significant differences are indicated with an asterisk (p < 0.05).

The response criterion (c) was not normally distributed, thus a one-tailed Wilcoxon test was calculated. Confirming our hypothesis, c was lower in the communicative (M = -0.10, SD = 0.52) than in the individual condition (M = -0.04, SD = 0.49) (Z = -1.66, p < .05, rank-biserial r = -0.31; Figure 2B). Additionally, we tested whether there was a response bias in the individual and communicative conditions towards ‘present’ or ‘absent’ responses, reflecting a positive or negative deviation from zero. While a one-sample Wilcoxon test revealed no significant response bias in the individual condition (Z = -0.81, p = 0.21, rank-biserial r = -0.15), responses in the communicative condition were significantly biased towards reporting the presence of a second agent (i.e. tendency to press ‘yes’) (Z = -1.73, p < 0.05, rank-biserial r = -0.32, one-tailed).

To analyze reaction times, a 2 × 2 ANOVA was calculated with the factors communicative/individual conditions and signal/noise trials. These were normally distributed. Confirming our hypothesis, responses occurred significantly faster in the communicative (M = 1601.42 ms, SD = 193.25 ms) than in the individual condition (M = 1622.90 ms, SD = 183.99 ms) (F(1,38) = 5.91, p < 0.05, partial η^2^ = 0.14) and faster for signal (M = 1586.12 ms, SD = 169.41 ms) than for noise trials (M = 1637.88 ms, SD = 198.01 ms; F(1,38) = 4.28, p < 0.05, partial η^2^ = 0.10 (Figure 2C). There was no interaction of experimental factors (F(1,38) = 0.08, p = 0.76, partial η^2^ = 0.00).

### Source-level Amplitude Results

#### Amplitude Results of Time Segment (1) Before Agent B’s Onset

In this time segment, only the contrast of the communicative versus individual condition was computed (see Table 1, red column). As this time segment is before agent B’s onset, participants cannot evaluate whether the trial’s outcome is non-expected/expected or whether agent B is present/absent.

**Table 1.**
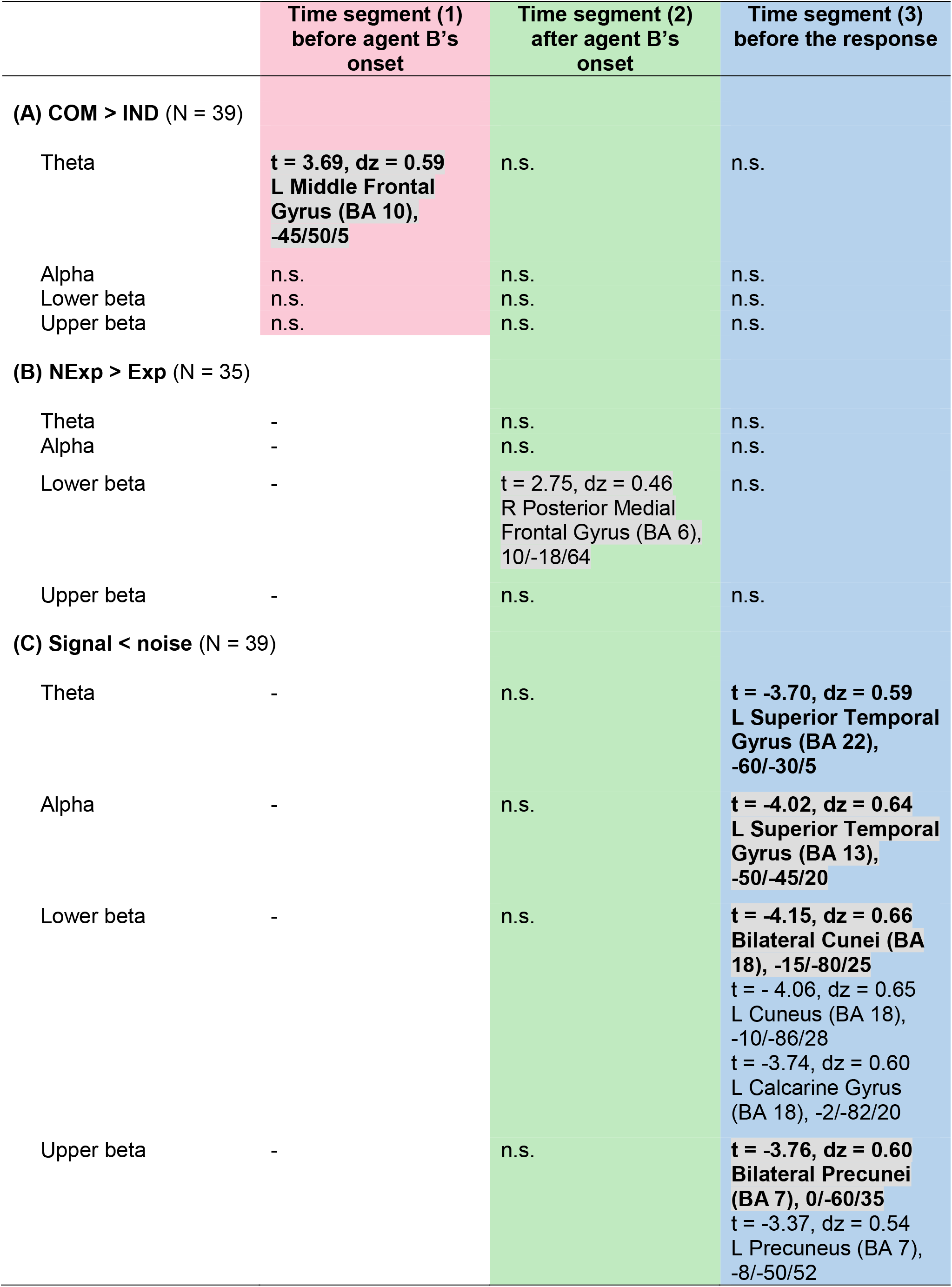
Statistical results of the EEG amplitude analyses. The columns depict the three time segments and the rows depict the contrasts divided into five frequency bands. The red, green and blue color indicate which contrasts were calculated for the time segments. In time segment (1) before agent B’s onset (i.e., observation of agent A), it is yet impossible to differentiate between non-expected and expected outcome or signal and noise trials (both dependent on agent B), thus only the contrast communicative (COM) versus individual (IND) condition (related to agent A) was computed for this time segment. The table indicates (A) a significant amplitude increase in the communicative (COM) in comparison to the individual (IND) condition (number of participants (N) = 39), (B) a significant amplitude increase in the non-expected (NExp) than expected (Exp) outcome (N = 35) and (C) a significant amplitude decrease in signal in comparison to noise trials (N = 39) in the indicated time segment, frequency band and brain area. These results are significant on the 5% alpha level (p < 0.05) and the t value, the effect size dz, the brain region, the Brodmann area (BA) and the coordinates according to the Montreal Neurological Institute (MNI) template (x/y/z) for the most significant voxel are indicated. The significant results from the *a priori* ROI-based analyses are written in normal black letters, the significant results from the data-driven whole brain analyses are written in bold and all results that are illustrated in Figure 3 are highlighted in grey. Non-significant effects are indicated with n.s..

##### A priori ROI-based Amplitude Analyses

The contrast of the communicative versus individual condition was computed for the 20 ROIs (see Supplementary Table S2), which showed a significant difference in the BOLD signal for the contrast communicative versus individual condition in the study of Zillekens et al. (2019). We could not find significant differences in the EEG amplitude analyses in the 20 ROIs in any frequency band in this time segment.

##### Data-Driven Whole Brain Amplitude Analyses

In the time segment (1) before agent B’s onset, significantly higher theta (4-7 Hz) amplitude was shown in the communicative than individual condition in the left middle frontal gyrus (p < 0.05, see Table 1A in bold letters for statistical parameters and coordinates, Figure 3A and for more brain views Figure S1).

**Figure 3.**
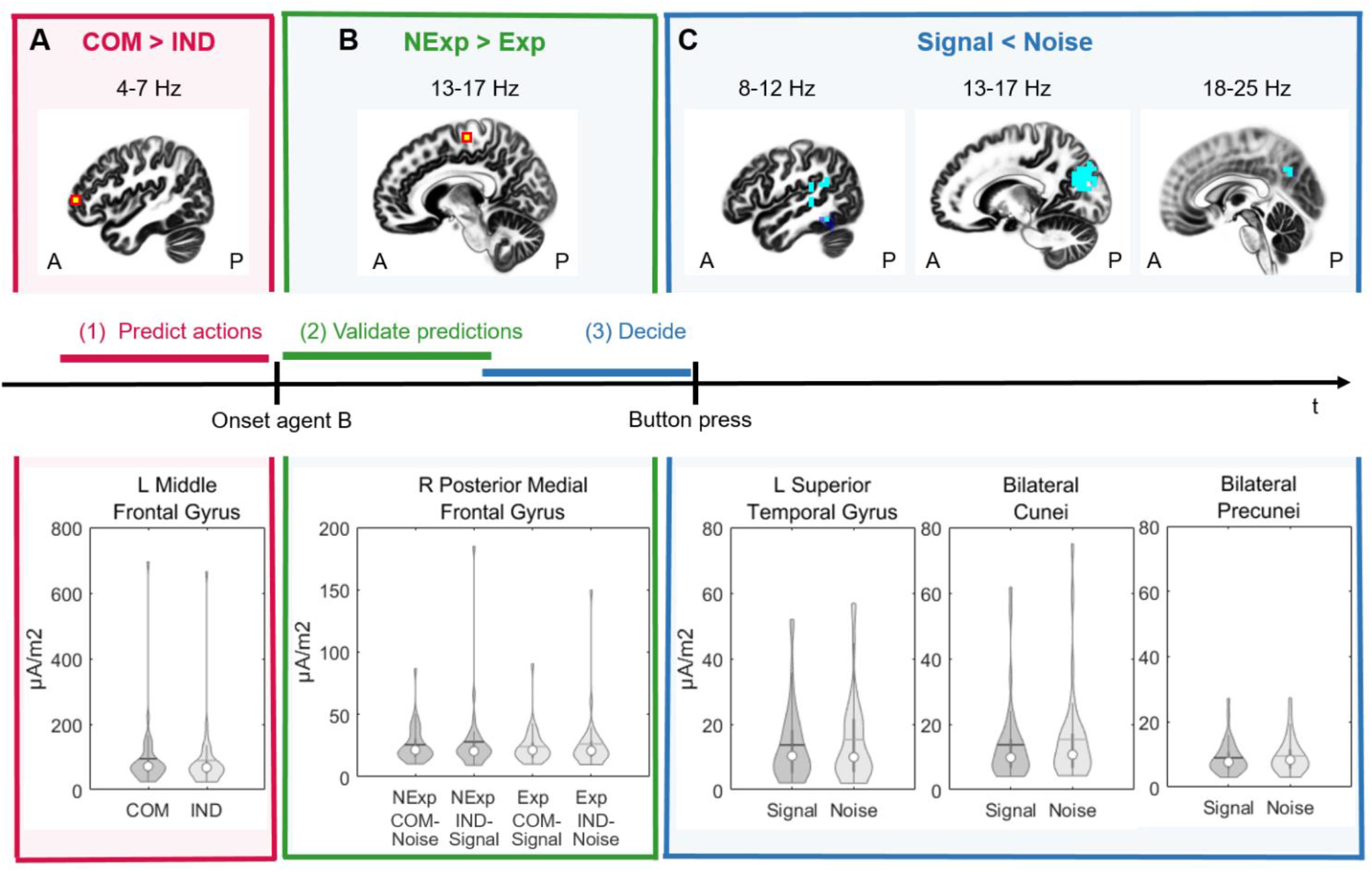
Brain patterns of the source-level amplitude changes. The top row depicts the voxels in the sagittal brain view, which show significant amplitude changes (A = anterior, P = posterior). Turquoise/blue colors indicate an amplitude decrease and yellow/red colors indicate an amplitude increase. The bottom row shows violin plots indicating the distribution of the data as well as the median (white circle) and mean (horizontal line) amplitude at the voxel with the most significant difference indicated in the brain maps above. (A) In the time segment (1) before agent B’s onset, higher amplitude was shown in the theta band in the communicative (COM) than in the individual (IND) condition in the left middle frontal gyrus (Brodmann area (BA) 10) with the peak difference at the MNI coordinates -45/50/5 (red section; Figure S1, S2). (B) In the time segment (2) after agent B’s onset, higher amplitude was shown in the lower beta band in the non-expected (NExp) than in the expected (exp) outcome in the right posterior medial frontal gyrus (BA 6) with the *a priori* defined MNI coordinates 10/-18/64 (Zillekens *et al*., 2019) (green section; Figure S3, S4). (C) In the time segment (3) before the response, lower amplitude was shown in the signal compared to the noise trials. This difference was significant in the left superior temporal gyrus in the theta (Figure S5A, S7A) but mainly in the alpha frequency band (left side of the blue section; Figure S5B, S7B). In the alpha frequency band, the most significant voxel was in the superior temporal gyrus in BA 13 (MNI: -50/-45/20) but there were also significant voxels in BA 41 in the superior temporal gyrus. Additionally, the middle temporal gyrus (BA 21 and 22) and the inferior temporal gyrus (BA 20), the supramarginal gyrus (temporal lobe, BA 40) as well as the inferior parietal lobule (parietal lobule, BA 40), the parahippocampal gyrus (BA 19, 27, 28, 30, 35 and 36), the lingual gyrus (BA 18, 19) and the fusiform gyrus (temporal and occipital lobe, BA 37) showed significantly lower alpha amplitude in the signal compared to noise trials. In the lower beta band, this difference showed peak significance in the bilateral cunei (BA 18; MNI: -15/-80/25; middle of the blue section; Figure S5C, S7C). This significant effect also spread to BA 7 and 19 of the cuneus (occipital lobe) and to the precuneus (parietal lobe, BA 31). In the *a priori* ROI-based analysis, the significant amplitude decrease in the signal compared to noise trials in the left cuneus (BA 18) was confirmed and additionally found in the left calcarine gyrus (BA 18) (please refer to Figure S6A, B). In the upper beta band, lower amplitude in the signal compared to noise trials was found in the bilateral precunei (parietal lobule, BA 7; MNI: 0/-60/35) (right of the blue section; Figure S5D, S7D). This significant difference spread slightly to the adjacent cingulate gyrus (BA 31) and was confirmed by the *a priori* ROI-based analysis (Figure S6C).

#### Amplitude Results of Time Segment (2) After Agent B’s Onset

In this time segment, all three contrasts (communicative versus individual condition, non-expected versus expected outcome and signal versus noise trials) were computed (see Table 1, green column). We could only find significant results in the contrast of the non-expected versus expected outcome:

##### A priori ROI-based Amplitude Analyses

The contrast of the non-expected versus expected outcome was tested within the 5 ROIs (see Supplementary Table S3), which showed a significant difference in the BOLD signal for the contrast non-expected (communicative noise trials and individual signal trials) versus expected (communicative signal trials and individual noise trials) outcome in the study of Zillekens et al. (2019). In the time segment (2) after agent B’s onset, significantly higher amplitude in the lower beta (13-17 Hz) frequency band (i.e., lower beta desynchronization/ lower activation) was shown in the non-expected compared to the expected outcome at the right posterior medial frontal gyrus (p < 0.05, see Table 1B written in normal black letters and Figure 3B).

##### Data-Driven Whole Brain Amplitude Analyses

No significant results were found.

#### Amplitude Results of Time Segment (3) Before the Response

In this time segment, again all three contrasts were computed (see Table 1, blue column). We could only find significant results in the contrast of signal versus noise trials:

##### A priori ROI-based Amplitude Analyses

The contrast of signal versus noise trials was computed for the 18 ROIs (see Supplementary Table S4), which showed a significant difference in the BOLD signal for the contrast signal versus noise trials in the study of Zillekens et al. (2019). In the time segment (3) before the response, the signal trials showed significantly lower beta amplitude (i.e., more desynchronization/ more activation) than the noise trials in the lower frequency range (13-17 Hz) at the left cuneus und the left calcarine gyrus and in the upper frequency range (18-25 Hz) at the left precuneus (*p* < 0.05, see Table 1C written in normal black letters).

##### Data-Driven Whole Brain Amplitude Analyses

In the time segment (3) before the response, the effect in the beta frequency band in the precuneus und cuneus was confirmed (p < 0.05, see Table 1C in bold letters and Figure 3C). Additionally, in the same time segment (3), the signal trials showed significantly lower amplitude in the theta and alpha frequency band at the left superior temporal gyrus (*p* < 0.05, see Table 1C and Figure 3C). In the alpha frequency band, the significant effect was stronger than in the theta band and spread to the middle temporal gyrus, the inferior temporal gyrus, the supramarginal gyrus, the inferior parietal lobule, the parahippocampal gyrus, the lingual gyrus and the fusiform gyrus (see Figure 3C and Figure S5 A and B).

### Summary of the Source-level and its Comparison to the Scalp-Level Amplitude Analyses

The results of the source-level amplitude analyses are illustrated in Figure 5. In time segment (1) before agent B’s onset, there was higher theta power in the communicative than individual condition in the left middle frontal gyrus. The coordinates of the middle frontal gyrus lie underneath the scalp-electrode AF7 (see Figure S2; Scrivener and Reader, 2022). Calculating the same contrast on scalp-level, we could confirm that in the time segment (1), power was significantly higher in the communicative than in the individual condition in the theta frequency band at AF7 (t = 1.79, dz = 0.29, p < 0.05, N = 39).

In time segment (2) after agent B’s onset, the source-level analyses revealed higher beta power in the non-expected than expected outcome in the posterior medial frontal gyrus. The coordinates correspond best to the scalp-level electrode C2 (see Figure S4; Scrivener and Reader, 2022). However, the significant source-level effect could not be confirmed on scalp-level (t = -0.73, p = n.s., N = 35).

In time segment (3) before the response, the source-level analyses revealed lower power in signal than in noise trials in the theta and alpha bands in the superior temporal gyrus and in the beta bands in the calcarine gyrus, cuneus and precuneus. All of these source-level effects could be replicated on scalp-level (see Figure S7; Scrivener and Reader, 2022): In this segment, power was significantly lower in signal than in noise trials in the theta frequency band at T7 (t = -2.72, dz = 0.44, p < 0.05, N = 39), in the alpha frequency band at TP7 (t = -3.53, dz = 0.57, p < 0.05, N = 39), in the lower beta frequency band at PO3 and POz (PO3: t = -3.65, dz = 0.58; PO_Z_: t = -2.85, dz =0.46; p < 0.05, N = 39).and in the upper beta frequency band at P_Z_ (t = -3.80, dz = 0.61, p < 0.05, N = 39).

### Exploratory Source-level Coherence Results

The exploratory coherence analyses were computed for all three time segments, but only for the contrast communicative versus individual condition. The reason for this choice is that it was based on the ROIs of Zillekens et al. (2019) and they only reported functional coupling differences between the communicative and individual condition. All coherence analyses were *a priori* ROI-based and computed for the 19 ROIs (see Supplementary Table S5), which showed a significant difference between the communicative and individual condition in the functional coupling to the amygdala in the study of Zillekens et al. (2019).

#### Coherence Results of Time Segment (1) Before Agent B’s Onset

In this time segment, a statistical trend emerged: We found marginally higher coherence in the communicative compared to the individual conditions in the alpha band (8-12 Hz) between the left superior orbital gyrus and the right anterior cingulate cortex (t = 4.20, p < 0.1, dz = 0.67, N = 39, Figure 4).

**Figure 4.**
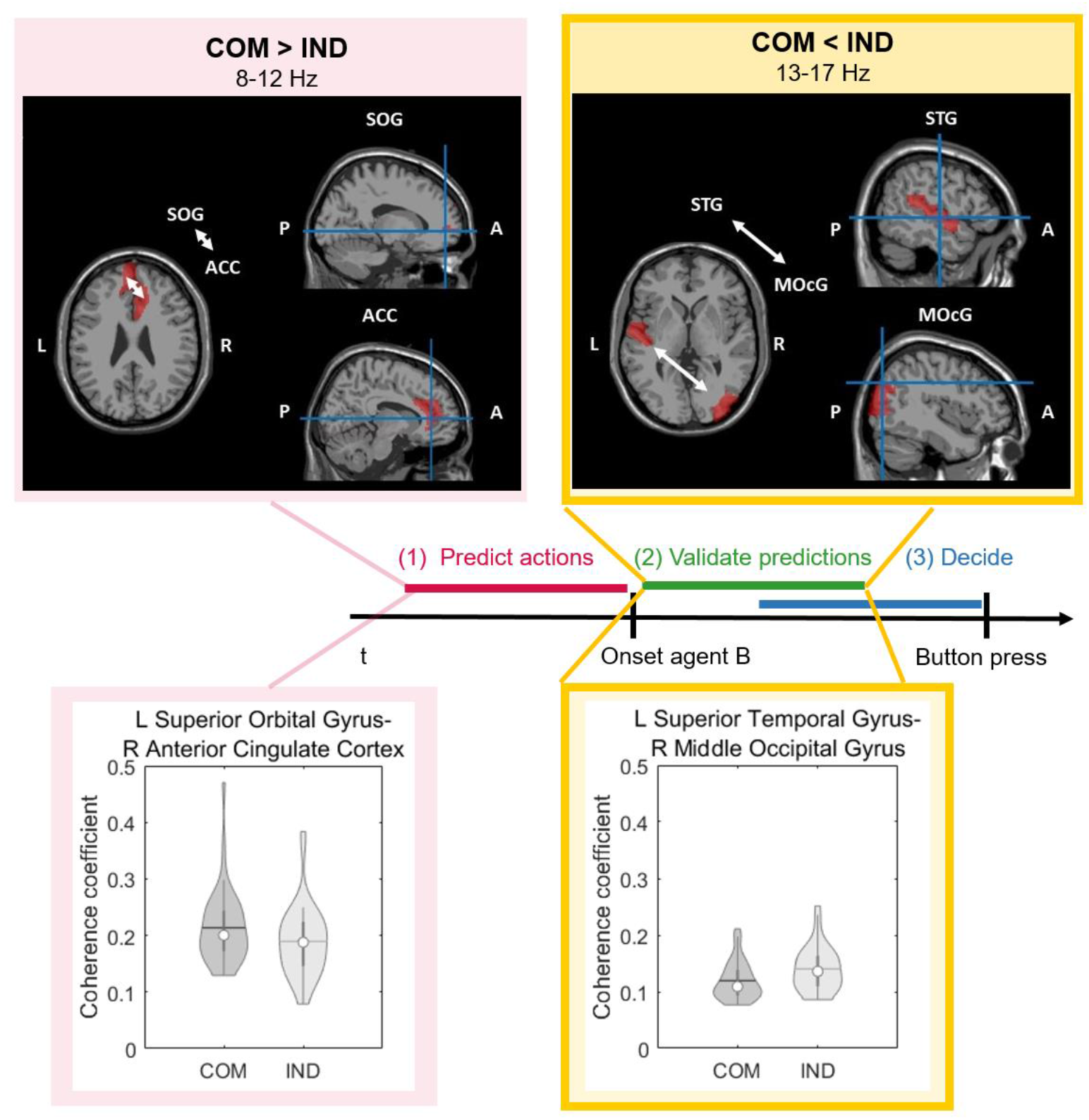
Brain patterns of the source-level coherence analyses. The frame around the yellow boxes indicate a significant result (p < 0.05), the missing frame around the red boxes indicate a statistical trend (p < 0.1). The top row depicts the brain regions (indicated in red), in which the MNI coordinates of the *a priori*-defined ROIs fall that showed significant lagged coherence. Slices on the left side of each panel are horizontal slices (L = left, R = right), those on the right side are sagittal slices (P = posterior, A = anterior) with the exact coordinates marked with a blue cross. The bottom row shows violin plots indicating the distribution of the data as well as the median (white circle) and mean (horizontal line) coherence coefficient between the two voxels indicated in the brain maps above. In the time segment (1) before agent B’s onset, higher alpha coherence between the left superior orbital gyrus (SOG; -16/52/-2) and the right anterior cingulate cortex (ACC; 10/38/6) was found in the communicative (COM) as compared to the individual (IND) condition by trend (p < 0.1; red sections). In the time segment (2) after agent B’s onset, significantly lower beta coherence was shown between the left superior temporal gyrus (STG; -52/-14/0) and the right middle occipital gyrus (MOcG; 42/-78/34) in the communicative (COM) as compared to the individual (IND) condition (p < 0.05; yellow/ green sections).

**Figure 5.**
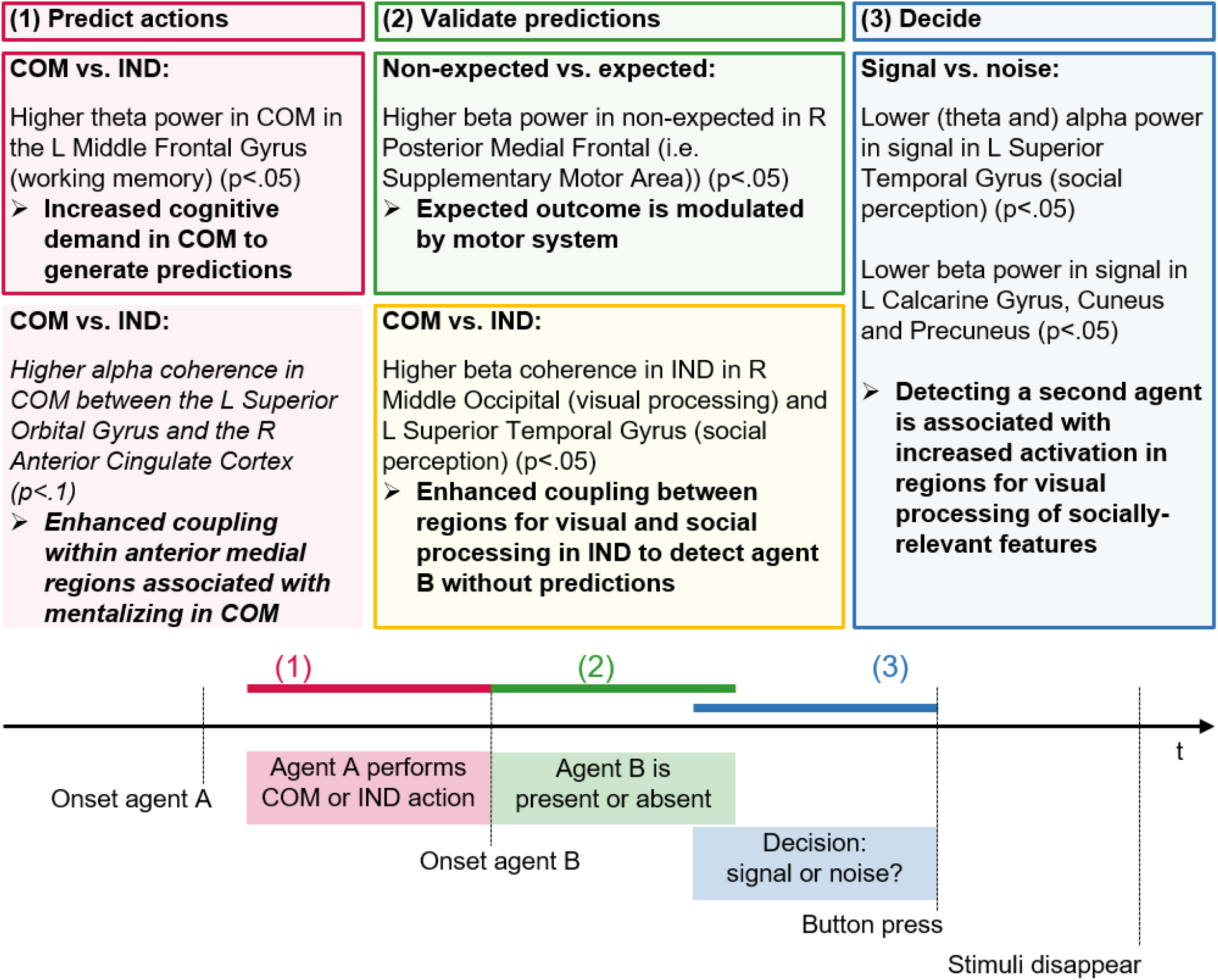
Spatio-temporal dynamics of source-level brain activity for interpersonal predictive coding. The results and their interpretation (in bold) are summarized in the text boxes for the time segments (1) before agent B’s onset (red sections), (2) after agent B’s onset (green and yellow sections) and (3) before the response (blue sections). The results from the amplitude analyses are indicated in the top row in the red, green and blue frames. The results from the coherence analyses for the communicative (COM) versus individual (IND) condition are indicated in the bottom row in red without a frame for time segment (1) and in yellow with a frame for time segment (2). The frames around the boxes indicate a significant result (p < 0.05), the missing frame around the red box in the bottom row and the italic font indicate a statistical trend (p < 0.1).

#### Coherence Results of Time Segment (2) After Agent B’s Onset

In the time segment (2) after agent B’s onset, we found significantly lower coherence in the communicative than the individual condition in the lower beta band (13-17 Hz) between the left superior temporal gyrus and the right middle occipital gyrus (t = -4.83, p < 0.05, dz = 0.77, N = 39, Figure 4).

#### Coherence Results of Time Segment (3) Before the Response

No significant results were found.

#### Summary of the Results of the Exploratory Source-Level Coherence Analyses

The results of the exploratory source-level coherence analyses are illustrated in Figure 5. Compared to the individual condition, coherence in the communicative condition was higher (statistical trend) in the alpha band between anterior medial regions in time segment (1) before agent B’s onset and significantly lower in the beta band between occipital and temporal regions in time segment (2) after agent B’s onset.

## DISCUSSION

The aim of this study was to investigate the spatio-temporal dynamics of oscillatory brain activity during interpersonal predictive coding, i.e. when we perceive and predict actions from nonverbal cues in point-light displays and use them to detect the presence of a second agent to whom the behavior has been directed. We expected different brain networks to be involved during the formation of predictions, the validation of the predictions and the actual detection of the second agent.

Our source-level results confirmed that different brain regions were active at different stages of the task (Figure 5). In time segment (1) before agent B’s onset, the communicative condition showed higher theta power and higher alpha coherence in frontal areas than the individual condition. In time segment (2) after agent B’s onset, the individual condition showed higher coherence between occipital and temporal regions compared to the communicative one. Additionally, we found higher beta power over the central brain area in the non-expected than expected outcome in this time segment. In time segment (3) before the response, signal trials demonstrated lower power than noise trials in lower frequencies in the temporal and in beta bands in the parietal and occipital regions. The source-level amplitude effects in the time segments (1) and (3) could be replicated on scalp-level, whereas the source-level effect in the time segment (2) remained non-significant on scalp-level.

The age, gender and AQ distribution of our sample was comparable to the preceding fMRI study by Zillekens et al. (2019). Using the same experimental design, the behavioral performance was similar across Zillekens’ and the present study.

Confirming our hypothesis, participants were more biased towards reporting the presence of a second agent (i.e., adopted a less conservative response strategy) after observing a communicative as compared to an individual gesture of a first agent (Manera, del Giudice, *et al*., 2011; Zillekens *et al*., 2019). In contrast to our hypotheses, the finding of increased sensitivity remained statistically non-significant when compared between the communicative and individual condition. Interestingly, Zillekens et al. (2019) could show a significant difference in in the sensitivity between the communicative and individual condition in their sample of the behavioral experiment, but this difference also remained non-significant in their fMRI experiment. One explanation for these inconsistent results is that the sensitivity effect was established with a slightly different experimental design: Manera and colleagues used a two-alternative forced-choice design (Manera, Becchio, et al., 2011). Participants were not asked to decide in every trial whether agent B was present or absent (as in our experimental design), but they always watched two trials and had to decide in which of the two trials agent B was present. That means that agent B was always present in one of the two trials. They found that the expectation of agent B after a communicative gesture led to better detectability of it. In contrast, in our experimental design, agent B was present only in 50% of the trials, which needed a decision (Manera, del Giudice, et al., 2011; Zillekens et al., 2019; Friedrich et al., 2022). In this case, the communicative gesture might have still led to better detectability of agent B (i.e., more hits) but at the same also led to seeing the Bayesian ghost (i.e., more false alarms). The Bayesian ghost is defined by a false alarm in the communicative noise trial (Friedrich et al., 2022). Based on agent A’s communicative gesture, participants expected agent B to be present in the cluster of noise dots and thus had an illusion of agent B although it was absent. More hits and more false alarms were reflected in a more liberal criterion but not in a higher sensitivity as shown in this study.

Concerning the reaction time, we could confirm our hypotheses that responses occurred faster in the communicative than in the individual condition and faster for signal than for noise trials.

On the neural level, we found different brain regions to be active during different stages of the social interaction task (see Figure 5):

For the time segment (1) before agent B’s onset, the source-level data-driven whole brain analysis revealed that the communicative condition showed significantly higher theta amplitude than the individual condition in the left middle frontal gyrus (Table 1, Figure 3 and 5). The coordinates fall into the left dorsolateral prefrontal cortex area of the gyrus and the effect could also be confirmed on scalp-level (Figure S1 and S2). A theta increase in the dorsolateral prefrontal cortex has been associated with working memory control processes (Braver *et al*., 1997; Wager and Smith, 2003; Sauseng *et al*., 2004; Sauseng *et al*., 2010; Barbey *et al*., 2013). This might indicate that the communicative condition recruited more working memory resources than the individual one as participants were forming predictions for agent B’s response action.

In contrast to our hypotheses, our EEG amplitude analyses did not show higher activation in the communicative compared to the individual condition of regions engaged in mirror neuron activity and mentalizing. Neither was increased activation demonstrated in any temporal (Isik *et al*., 2017; Walbrin *et al*., 2018) or prefrontal networks previously associated with social interaction processing (Sliwa and Freiwald, 2017). Zillekens et al. (2019) did not find activation of these regions when contrasting the communicative versus individual conditions either. One explanation is that both conditions were perceived in terms of social biological motion as they included point-light displays showing moving agents (Zillekens *et al*., 2019). In another recent publication from our lab, we have demonstrated higher sensorimotor activation in time segment (1) in false alarm compared to correct rejection and hit trials within the communicative condition (Friedrich *et al*., 2022). We had interpreted this sensorimotor activation as a neural signature of generating predictions that outweigh sensory information presented later in the time segment (3) and thus leading to the illusion of seeing agent B (i.e., to seeing a Bayesian ghost). The fact that we did not find a difference in sensorimotor activation between the individual and communicative condition in the current data indicates that this sensorimotor activation is not related to generating predictions generally (which is possible in the communicative but not in the individual condition), but rather to generating top-down predictions that outweigh the bottom-up sensory information and thus lead to the perception of a Bayesian ghost.

In the exploratory source-level coherence analyses a statistical trend emerged for the contrast of the communicative versus individual condition, which corroborates the results of Zillekens et al. (2019). Zillekens and colleagues (2019) had found that the amygdala was functionally coupled to the medial prefrontal cortex in communicative trials, whereas it increased its functional connectivity with fronto-parietal areas in the individual condition.

Here, EEG coherence analyses between the cortical regions (note, that in the EEG no signal from the amygdala could be captured) revealed higher alpha coherence between superior orbital gyrus and the anterior cingulate cortex in the communicative compared to the individual condition during the observation of agent A (Figure 4 and 5). Both of these medial anterior regions are important for social tasks and might be involved in mentalizing processes (Amodio and Frith, 2006; Monticelli *et al*., 2021). This means that the observation of communicative gestures and the resulting inference of actions in the time segment before the onset of agent B, may not only have led to the recruitment of more cognitive resources, but also to enhanced coupling and thus to a more efficient network within medial anterior regions associated with social and mentalizing processes.

In the time segment (2) after agent B’s onset, participants could validate their predictions and try to detect agent B in a cluster of moving dots. Indicating the higher difficulty level of detecting a second agent without expectations about the specific movement, enhanced coupling between the middle occipital and the superior temporal gyrus was shown in the individual compared to the communicative condition in our source-level coherence analysis (Figure 4 and 5). It has been shown that the visual system is sensitive to point-light biological motion (Pavlova, 2012). The superior temporal gyrus is a region involved in social perception and cognition (Baron-Cohen *et al*., 1999; Jou *et al*., 2010). The superior temporal gyrus is strongly connected to the adjacent posterior superior temporal sulcus, a brain region part of the mirror neuron system and mentalizing system (Rizzolatti *et al*., 2001; Schilbach *et al*., 2013; Yang *et al*., 2015). Thus, it has been associated with biological motion perception and social interaction (Caspers *et al*., 2010; Pavlova, 2012; Isik *et al*., 2017; Walbrin *et al*., 2018). Moreover, Quadflieg and colleagues (2015) showed that both the left and right posterior superior temporal sulcus were more active in incongruent interactions (i.e., two persons performing non-matching individual actions) in comparison to congruent ones (i.e., two people performing a communicative action) (Quadflieg *et al*., 2015), which is in line with our results.

In the time segment (2) after agent B’s onset, we had furthermore assumed that the better predictability of an expected outcome leads to increased motor-relevant cortex activation as compared to a non-expected outcome (Braukmann *et al*., 2017). Indeed, our source-level amplitude analyses showed lower beta amplitude (i.e., beta suppression) in the expected outcome in comparison to the non-expected outcome after agent B’s onset in the right posterior medial frontal gyrus (Table 1, Figure 3 and 5). The posterior medial frontal gyrus includes Brodmann Area 6, which consists of the premotor and supplementary motor area. The coordinates of the ROI in our analysis fall into the supplementary motor area (Tzourio-Mazoyer *et al*., 2002; Kim *et al*., 2010) (see Figure 3 and Figure S3). As beta suppression over motor areas reflects motor-relevant activation (McFarland *et al*., 2000), our source-level amplitude results show evidence in favor of our hypothesis. However, this effect remained non-significant on scalp-level (Figure S4). Differences in scalp- and source-level analyses over the motor area might arise from the folding of the underlying cortex.

Depending on the specific folding of the central sulcus and the direction of the dipoles, the signal at the source might be projected to different scalp-locations and not only to the electrode lying directly above the source, and there might be considerable spatial smearing on scalp-level. That is why we think that the source-level results are a better indicator of the true signal than the scalp-level analyses. Still this reported finding should be further confirmed in future studies.

Also because in contrast to our source-level results, Zillekens et al. (2019) reported a higher BOLD response (i.e., more activation) for the non-expected than the expected outcome in the right posterior medial frontal gyrus together with the left cerebellum, left fusiform gyrus and parietal areas. This effect has been interpreted as being associated with the computation of error signals and the different processing of incongruent and congruent stimuli (Zillekens *et al*., 2019). It seems that this motor-relevant brain area can be activated in both cases: first, during the processing of expected actions (i.e., predictable actions and normal goal-directed movements; Tzagarakis *et al*., 2010; Braukmann *et al*., 2017; Cheng *et al*., 2017) and second, during the processing of errors (i.e., erroneous or distorted movements; Meyer *et al*., 2016; Cheng *et al*., 2017).

In the time segment (3) before the response, the decision had to be made whether to press ‘yes’ or ‘no’ and thus indicate whether agent B was present or absent. The number of dots displayed were kept stable over signal and noise trials. While we found a significant effect of signal versus noise trials in our source-level amplitude analyses (Table 1, Figure 3, 5 and S5), this difference did not become significant in the motor system as hypothesized. We found more activation (i.e., lower alpha and beta amplitude) in signal than noise trials in the occipital lobe, in the cuneus, the calcarine sulcus and the lingual gyrus which are involved in visual processing (Flores, 2002). Additionally, the temporal lobe was more activated in signal than noise trials: The superior temporal gyrus showed a strong effect in the alpha band with a marginal effect also in the theta band, which was probably driven by alpha smearing into lower frequency bands. As already mentioned above, the superior temporal gyrus is a region involved in social perception and cognition (Baron-Cohen *et al*., 1999; Jou *et al*., 2010) and its adjacent posterior superior temporal sulcus is associated with social interaction processing (Isik *et al*., 2017; Walbrin *et al*., 2018).

Moreover, we found more alpha decrease (i.e., more activation) in the middle temporal gyrus, the inferior temporal gyrus, the fusiform gyrus and the parahippocampal gyrus in signal than noise trials. The temporal gyrus is part of the ventral stream of visual processing and involved in object recognition (Kravitz *et al*., 2014). The fusiform gyrus is specific to face recognition (Kanwisher *et al*., 1997), whereas the parahippocampal gyrus is engaged in visual recognition of scenes (Epstein and Kanwisher, 1998).

The fact that the differences in the occipitotemporal regions remained non-significant between the communicative and individual condition but became significant for the comparison of signal and noise trials in this time segment before the response might be explained by the recent work of Wurm and Caramazza (2019). They suggested that the activation of the occipitotemporal regions is rather associated with visual processing of socially-relevant features such as the presence of a person, the directedness of actions or orientation of agents than an abstract representation of sociality (Wurm and Caramazza, 2019). In the present study, it seems that these regions are more sensitive to whether there is actually a second agent that can be perceived (i.e., presence/absence of the second agent) than whether the two agents interact with each other or not (communicative versus individual condition). These results also highlight the importance to look at different time segments and thus entangle the different cognitive processes involved in social tasks.

In the parietal lobe, we found more alpha decrease (i.e., more activation) in signal than noise trials in Brodman Area 40 of the supramarginal gyrus and the inferior parietal lobule. In the precuneus, there was an amplitude decrease in signal compared to noise trials in the beta band in the present study. While the Brodmann Area 40 was shown to be activated during mirror-like processing of action observation (Del Vecchio *et al*., 2020), the precuneus is part of the mentalizing system (Van Overwalle and Baetens, 2009). Among others, the precuneus is involved in motor and mental imagery, processing of visuo-spatial information and attention as well as processing and understanding intentions and actions (Cavanna and Trimble, 2006; Schilbach *et al*., 2006; Schilbach *et al*., 2008; Schilbach *et al*., 2012; Brandi *et al*., 2021). As the effects in alpha and beta bands in the precuneus, cuneus and calcarine gyrus did not replicate the results from Zillekens et al. (2019), the results in the present study might represent a very specific activation right before the response was given. Contrary to analyzing the activation over the entire video duration, the strength of the current approach using EEG was to disentangle the specific cognitive processes involved in this complex task.

Taken together, in the time segment (3) before the response, our source-level analyses showed that signal compared to noise trials led to increased activation in regions relevant for visual processing, social perception and cognition. These results could also be confirmed on scalp-level (Figure S7). Signal and noise trials did not differ significantly in the predicted motor areas. It seems that activation of the sensorimotor cortex is specific to perceiving a biological motion (or illusion) after a communicative gesture (Friedrich *et al*., 2022). In the present study, we contrasted correct signal (i.e., hit) versus correct noise (i.e., correct rejection) trials collapsed over the communicative and individual condition. Our findings suggest that detecting a second agent correctly (i.e., providing the basis for a social interaction independently whether it follows a communicative or individual gesture) is associated with increased activation of regions involved in visual processing of socially-relevant features.

### Limitations

We based this study on Zillekens et al. (2019) in order be able to perform source-level EEG analyses based on *a priori* ROIs that have been found to be important for interpersonal predictive coding using fMRI. This way, we aimed to combine the advantage of the fMRI’s high spatial resolution with the EEG’s high temporal resolution to investigate spatio-temporal dynamics of interpersonal predictive coding. One limitation of our study is, however, that we did not increase the EEG’s spatial resolution in order to more accurately detect the sources by using high-density EEG caps combined with individual MRI scans or by localizing the electrodes on the scalp individually in the three dimensional space. We still believe that we can differentiate between the fMRI-based *a priori* ROIs as it was shown that an exact source reconstruction based on 64 scalp channels is possible using LORETA (Akdeniz, 2016). The improvement in source localization accuracy is plateauing after the use of about 63 channels (Lantz et al., 2003; Sohrabpour et al., 2015) and 60 channels might even be more robust against higher noise levels than a higher number of electrodes (Chauveau et al., 2008).

Although this study was based on the results of Zillekens *et al*. (2019), we performed the hypotheses-driven amplitude analyses not only on their source-level *a prior* defined ROIs but also on scalp-level electrodes and based on a data-driven approach. We investigated differences in amplitude over different time segments, contrasts and frequency bands. Additionally, we performed exploratory coherence analyses. Due to this rather exploratory nature of our study -and like most novel results-, it will be beneficial to replicate these results in future studies.

The division of the data in the three time segments also has its limitation: The time segment (1) before agent B’s onset only included the last second of agent A’s action (i.e., on average the first 400 ms were not analyzed) and the time segments (2) and (3) were overlapping (on average about 400 ms) in our analyses. Thus, we were not able to capture the whole process of forming predictions and the process of validating the predictions and deciding the response could not be disentangled entirely. The reason for this is that we needed time segments with the same length to keep the number of samples and frequency resolution equal across analyses. If we increased the duration of the time segments in order to analyze the whole action of agent A in time segment (1), time segments (2) and (3) would overlap even more. And if we decreased the duration of the time segments in order to have less overlap between time segments (2) and (3), we would lose more time of agent A’s action in time segment (1).

Despite the rather exploratory nature of our study and the limitations with the division of time segments, our results provide valuable insights about which brain areas are involved in the different time segments during interpersonal predictive coding. The mere fact that we find significant differences in the contrasts specifically for the hypothesized time segments (i.e., first differences between the communicative and individual condition, then differences between expected and non-expected outcomes and last differences between signal and noise trials) demonstrate that our analyses were able to reliably capture and disentangle the different processes.

## CONCLUSION

In this study, we demonstrated the neural spatio-temporal dynamics of perceiving and predicting actions from nonverbal social cues and of using them to optimize response behavior by means of EEG. First, the perception of communicative actions and the resulting predictions led to an enhanced functional coupling within medial anterior regions involved in social and mentalizing processes and a higher deployment of cognitive resources than required in the individual condition. Second, the detection of a social agent without the help of action predictions, led to enhanced coupling between regions for social perception and visual processing in the individual condition. In line with our hypothesis, an expected outcome was modulated by activation of the motor system. Third, the correct detection of a second agent was associated with increased activation in areas for visual processing of socially-relevant features rather than the predicted motor areas. Overall, our results confirm that different neural markers associated with different cognitive processes are dominant at different stages of social interactions and that it is crucial to consider the temporal dynamics during social interactions and their neural correlates.

Future studies could be extended to include individuals with social interaction difficulties or ‘disorders of social interaction’ (Schilbach, 2016) to investigate whether they show aberrant brain activity in the dynamic brain patterns described in this study. This would allow conclusions about which processes (cognitive, motor, visual or social) and stages (formation of predictions, validation of predictions, decision processes) are affected and may contribute to interactional problems. Here, an interesting avenue could be the use of brain stimulation techniques or neurofeedback training (Pineda *et al*., 2008; Pineda *et al*., 2014; Friedrich *et al*., 2015; Nitsche and Paulus, 2011; Herrmann *et al*., 2013; Amatachaya, 2014), to improve social interaction.

## Acknowledgements

We would like to thank Cristina Becchio and Atesh Koul for providing the experimental task and advice during setup and analysis.

This study was funded by grants from the Deutsche Forschungsgemeinschaft (DFG) to EVCF (FR 3961/1-1) and to PS (SA 1782/2-2) and by the Max Planck Society via a grant for an independent Max Planck Research Group to LS.

## Authors contributions

Conceptualization, E.V.C.F., I.C.Z., L.S., and P.S.; Methodology, E.V.C.F., A.L.B., I.C.Z., and P.S.; Software, E.V.C.F., A.L.B., and I.C.Z.; Validation, E.V.C.F., I.C.Z., A.L.B., and P.S.; Formal Analysis, E.V.C.F., I.C.Z., A.L.B., D.O., E.V.S., and J.S.; Investigation, E.V.C.F., D.O., and E.V.S.; Data Curation, E.V.C.F., A.L.B., D.O., E.V.S., and J.S.; Resources, P.S.; Writing – Original Draft, E.V.C.F., D.O., L.S., and P.S.; Writing – Review & Editing, E.V.C.F., I.C.Z., A.L.B., D.O., E.V.S., J.S., L.S., and P.S.; Visualization, E.V.C.F. J.S.; Supervision, L.S. and P.S.; Project Administration, E.V.C.F., L.S., and P.S.; Funding Acquisition, E.V.C.F., L.S., and P.S.

## List of abbreviations

A: anterior
ACC: anterior cingulate cortex
AQ: Autism Spectrum Quotient
BA: Brodmann area
BOLD: blood oxygenation level dependent
c: response criterion
COM: communicative
d’: sensitivity
EEG: electroencephalography
EOG: electrooculogram
Exp: expected
FA: false alarm
fMRI: functional magnetic resonance imaging
IND: individual
L: left
M: mean
MNI: Montreal Neurological Institute
MOcG: middle occipital gyrus
NExp: non-expected
P: posterior
R: right
ROI(s): region(s) of interest
SD: standard deviation
sLORETA: standardized Low Resolution Electromagnetic Tomography
SOG: superior orbital gyrus
STG: superior temporal gyrus

