## Supplementary Material for "Spatio-Temporal Dynamics of Oscillatory Brain Activity during the Observation of Actions and Interactions between Point-light Agents"

**Table S1.** *Number of trials for the EEG analyses.* **(A)** For the EEG amplitude analyses, for the contrasts communicative (COM) versus individual (IND) conditions, signal versus noise trials and non-expected (COM-Noise, IND-Signal) versus expected (COM-Signal, IND-Noise) outcomes, the number of included participants (N) and the mean, standard deviation (SD), minimum number (min) and maximum number (max) of the trials used per condition are displayed. Due to the artifact rejection, it is possible that the number of trials differed between time segments. Time segments (1) before and (2) after agent B's onset were extracted together from the continuous EEG data, whereas time segment (3) before the response was extracted separately from the other time segments. **(B)** For the EEG coherence analyses, the same information about the number of trials is displayed for the contrast communicative (COM) versus individual (IND) conditions.

| | | **Time segments (1) and (2)** | | | | **Time segment (3)** | | | |
|---|---|---|---|---|---|---|---|---|---|
| | N | Mean | SD | Min | Max | Mean | SD | Min | Max |
| **(A) Amplitude** | | | | | | | | | |
| COM | 39 | 77.44 | 15.00 | 43 | 99 | 79.69 | 14.54 | 46 | 101 |
| IND | 39 | 77.85 | 15.55 | 46 | 107 | 80.62 | 15.49 | 50 | 114 |
| Signal | 39 | 83.03 | 22.28 | 24 | 121 | 86.15 | 23.31 | 27 | 123 |
| Noise | 39 | 72.26 | 21.32 | 29 | 120 | 74.15 | 20.41 | 32 | 119 |
| COM-Noise | 35 | 35.80 | 10.01 | 22 | 59 | 36.63 | 9.66 | 23 | 59 |
| IND-Signal | 35 | 41.74 | 9.52 | 20 | 58 | 44.94 | 14.94 | 21 | 108 |
| COM-Signal | 35 | 43.89 | 10.01 | 24 | 63 | 47.03 | 15.29 | 27 | 114 |
| IND-Noise | 35 | 37.89 | 10.74 | 20 | 61 | 38.97 | 10.21 | 21 | 60 |
| **(B) Coherence** | | | | | | | | | |
| COM | 39 | 74.85 | 14.92 | 43 | 99 | 77.28 | 14.73 | 46 | 101 |
| IND | 39 | 74.85 | 14.92 | 43 | 99 | 77.28 | 14.73 | 46 | 101 |

**Table S2** (adapted from the top part of the Supplementary Table S5 in Zillekens et al. (2019)). *ROIs for the effect of the communicative (COM) versus individual (IND) condition*. The presented ROIs showed significant differences in the BOLD signal in the contrasts **(A)** COM > IND and **(B)** IND > COM in the fMRI study from Zillekens et al. (2019). Anatomical labels and coordinates according to the Montreal Neurological Institute (MNI) template are displayed for the regions that could be recorded with EEG and were tested in the current study.

| **Macroanatomical location** | **x** | **y** | **z** |
|---|---|---|---|
| **(A) COM > IND** | | | |
| R Superior Frontal Gyrus | 22 | 56 | 24 |
| R Superior Medial Gyrus | 30 | 62 | 8 |
| R Middle Orbital Gyrus | 42 | 46 | 0 |
| **(B) IND > COM** | | | |
| L Superior Parietal Lobule | -34 | -70 | 54 |
| L Inferior Parietal Lobule | -36 | -50 | 56 |
| L Superior Occipital Gyrus | -20 | -66 | 30 |
| L Angular Gyrus | -44 | -72 | 40 |
| L Inferior Temporal Gyrus | -50 | -48 | -10 |
| L Inferior Occipital Gyrus | -46 | -78 | -12 |
| L Middle Temporal Gyrus | -66 | -54 | -6 |
| R Postcentral Gyrus | 40 | -28 | 40 |
| R Precentral Gyrus | 26 | -28 | 72 |
| R Superior Frontal Gyrus | 38 | -12 | 64 |
| R Middle Frontal Gyrus | 36 | -6 | 64 |
| R Inferior Occipital Gyrus | 40 | -70 | -6 |
| R Inferior Temporal Gyrus | 46 | -50 | -14 |
| L IFG (p. Opercularis) | -38 | 4 | 30 |
| L IFG (p. Triangularis) | -52 | 14 | 36 |
| L Precentral Gyrus | -54 | 8 | 36 |
| L Middle Frontal Gyrus | -44 | 24 | 42 |

**Table S3** (adapted from the Supplementary Table S6 in Zillekens et al. (2019)). *ROIs for the effect of expected versus non-expected outcome*. The presented ROIs showed significant differences in the BOLD signal in **(A)** a contrast of expected outcome [(COM_Signal + IND_Noise) > (IND_Signal + COM_Noise)] and **(B)** a reversed contrast of non-expected outcome [(COM_Noise + IND_Signal) > (COM_Signal + IND_Noise)] in the fMRI study from Zillekens et al. (2019). Anatomical labels and MNI coordinates are displayed for the regions that could be recorded with EEG and were tested in the current study.

| Macroanatomical location | x | y | z |
|---|---|---|---|
| **(A) Expected outcome** | | | |
| No suprathreshold cluster | | | |
| **(B) Non-expected outcome** | | | |
| R Posterior-Medial Frontal | 10 | -18 | 64 |
| R Precuneus | 10 | -50 | 62 |
| L Paracentral Lobule | -8 | -36 | 60 |
| L Precuneus | -10 | -54 | 62 |
| L Inferior Temporal Gyrus | -44 | -44 | -22 |

**Table S4** (adapted from the bottom part of the Supplementary Table S5 in Zillekens et al. (2019)). *ROIs for the effect of signal versus noise trials*. The presented ROIs showed significant differences in the BOLD signal in the contrasts **(A)** Signal > Noise and **(B)** Noise > Signal in the fMRI study of Zillekens et al. (2019). Anatomical labels and MNI coordinates are displayed for the regions that could be recorded with EEG and were tested in the current study.

| **Macroanatomical location** | **x** | **y** | **z** |
|---|---|---|---|
| **(A) Signal > Noise** | | | |
| L Anterior Cingulate Cortex | -2 | 36 | 20 |
| **(B) Noise > Signal** | | | |
| L Cuneus | -10 | -86 | 28 |
| L Calcarine Gyrus | -2 | -82 | 20 |
| R Calcarine Gyrus | 20 | -98 | 8 |
| R Temporal Pole | 56 | 4 | 2 |
| R Middle Temporal Gyrus | 60 | -2 | -14 |
| R Superior Temporal Gyrus | 60 | -12 | 2 |
| R Precentral Gyrus | 52 | 0 | 34 |
| L Precuneus | -8 | -50 | 52 |
| R Precuneus | 2 | -62 | 50 |
| R Medial Cingulate Cortex | 6 | -42 | 52 |
| L Inferior Parietal Lobule | -26 | -52 | 60 |
| R Superior Frontal Gyrus | 30 | 60 | 26 |
| R Middle Frontal Gyrus | 28 | 56 | 32 |
| L Posterior-Medial Frontal | 0 | -2 | 54 |
| R Posterior-Medial Frontal | 10 | 4 | 52 |
| L Medial Cingulate Cortex | -12 | -10 | 52 |
| L Precentral Gyrus | -30 | -16 | 58 |

**Table S5** (adapted from the Supplementary Table S8 in Zillekens et al. (2019)). *ROIs resulting from a psychophysiological interaction analysis revealing peak clusters of co-activation with neural signaling in the left amygdala for the communicative (COM) and individual (IND) condition.* The table displays peak regions coupled to amygdala functioning in the context of **(A)** IND > COM and **(B)** COM > IND from the fMRI study from Zillekens et al. (2019). Anatomical labels and MNI coordinates are displayed for the regions that could be recorded with EEG and were tested in the current study.

| **Macroanatomical location** | **x** | **y** | **z** |
|---|---|---|---|
| **(A) IND > COM** | | | |
| L Inferior Temporal Gyrus | -58 | -42 | -16 |
| L Inferior Occipital Gyrus | -50 | -64 | -8 |
| R Superior Parietal Lobule | 32 | -58 | 64 |
| R Inferior Parietal Lobule | 54 | -44 | 52 |
| R SupraMarginal Gyrus | 46 | -32 | 36 |
| L Middle Frontal Gyrus | -42 | 38 | 22 |
| L Inferior Frontal Gyrus | -48 | 44 | 16 |
| R Middle Temporal Gyrus | 44 | -72 | 28 |
| R Middle Occipital Gyrus | 42 | -78 | 34 |
| R Middle Frontal Gyrus | 46 | 14 | 52 |
| R Inferior Temporal Gyrus | 58 | -60 | 2 |
| L Inferior Parietal Lobule | -36 | -66 | 56 |
| L Superior Parietal Lobule | -26 | -72 | 58 |
| **(B) COM > IND** | | | |
| L Superior Medial Gyrus | -2 | 50 | 6 |
| L Superior Orbital Gyrus | -16 | 52 | -2 |
| R Superior Medial Gyrus | 16 | 46 | 8 |
| R Anterior Cingulate Cortex | 10 | 38 | 6 |
| L Temporal Pole | -42 | 12 | -14 |
| L Superior Temporal Gyrus | -52 | -14 | 0 |

**Figure S1.** *Source-level amplitude effect based on the data-driven whole brain analyses for the contrast communicative versus individual condition.* In the time segment (1) before agent B's onset, power was significantly ($p < 0.05$) higher in the communicative than in the individual condition in the theta frequency band in the left middle frontal gyrus. Significant voxels are marked in yellow/red colors, the most significant voxel lies at the MNI coordinates -45/50/5.

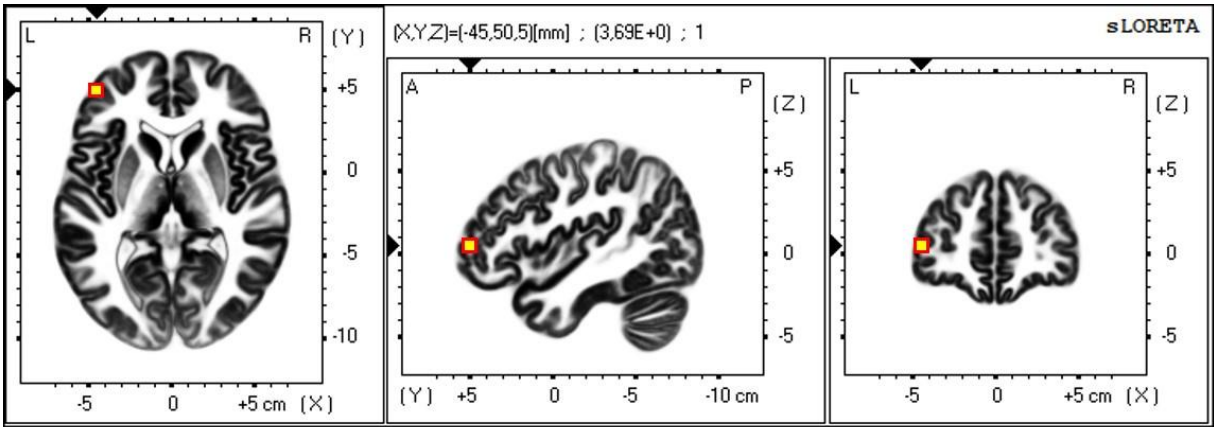

**Figure S2.** *Scalp-level amplitude effect for the contrast communicative versus individual condition.* The coordinates -45/50/5 of the left middle frontal gyrus (Figure S1) correspond best to the scalp-level electrode AF7 (see Table 1 in Scrivener and Reader (2022)). We calculated the same contrast with the scalp-level electrode AF7 (indicated with a red circle), which reached significance in the source-space for the left middle frontal gyrus (-45/50/5). In the time segment (1) before agent B's onset, power was significantly higher in the communicative than in the individual condition in the theta frequency band at AF7 ($p < 0.05$). The result is indicated with the yellow filling of electrode AF7 and is in line with the significant source-level effect illustrated in Figure S1.

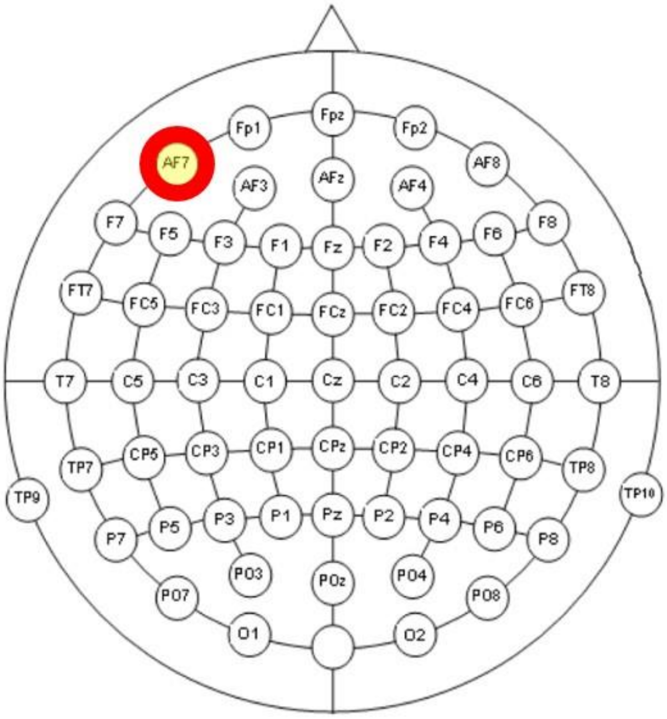

**Figure S3.** *Source-level amplitude effect based on the a priori ROI-based analyses for the contrast non-expected versus expected outcome.* In the time segment (2) after agent B's onset, power was significantly higher ($p<0.05$) in the lower beta band in the non-expected than in the expected outcome in the right posterior medial frontal gyrus. The exact voxel of the *a priori* ROI is marked in yellow/red colors and lies at the MNI coordinates 10/-18/64.

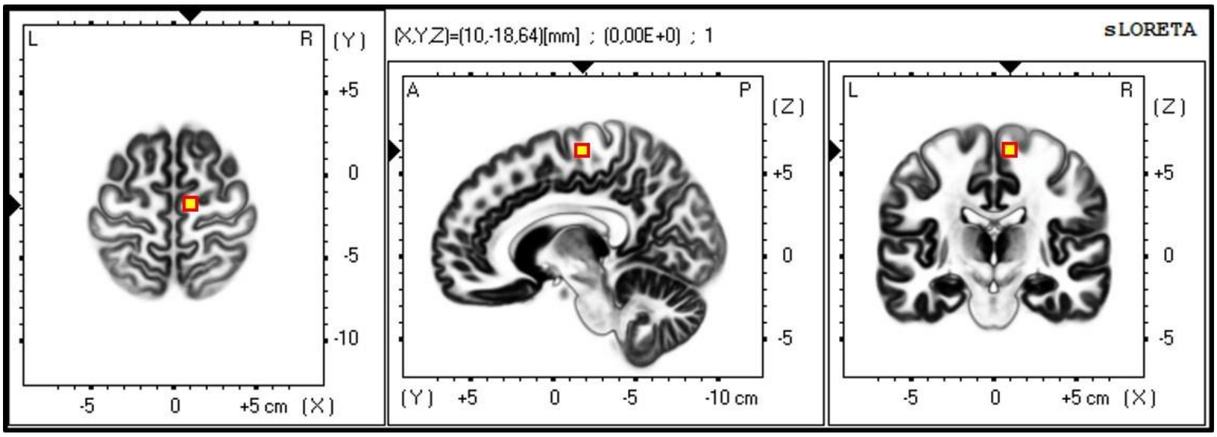

**Figure S4.** *Scalp-level amplitude effect for the contrast non-expected versus expected outcome.* The coordinates 10/-18/64 of the right posterior medial frontal gyrus (Figure S3) correspond best to the scalp-level electrode C2 (see Table 1 in Scrivener and Reader (2022)). We calculated the same contrast with the scalp-level electrode C2 (indicated with a red circle), which reached significance in the source-space for the right posterior medial frontal gyrus (10/-18/64.; see Figure S3). There was no significant difference between the non-expected and expected outcome in the lower beta band in the time segment (2) after agent B's onset (p = n.s.; no filling).

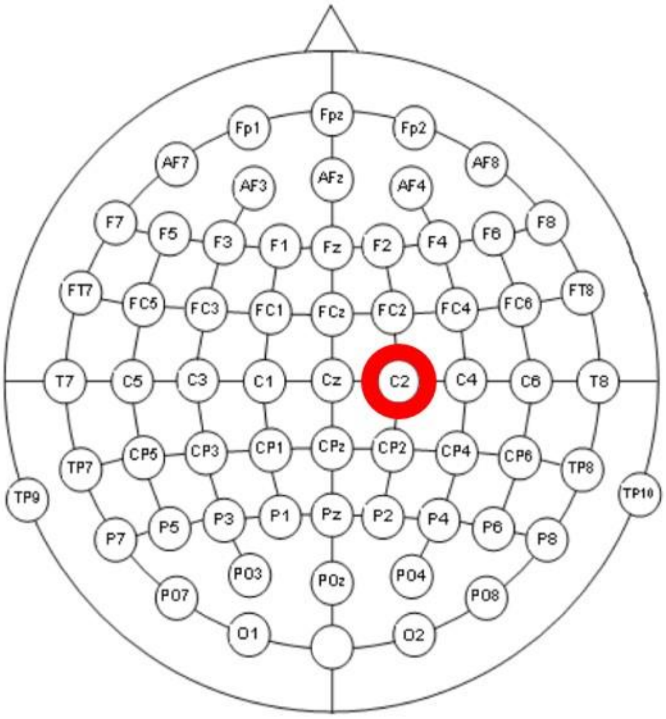

**Figure S5.** *Source-level amplitude effect based on the data-driven whole brain analyses for the contrast signal versus noise trials*. Significant voxels are marked in turquoise/blue colors in the figure and the MNI coordinates for the most significant voxel is indicated in brackets in the text. In the time segment (3) before the response, power was significantly ($p < 0.05$) lower in signal than in noise trials **(A)** in the theta frequency band (blue letter) in the left superior temporal gyrus (-60/-30/5) **(B)** in the alpha band (green letter) in the left superior temporal gyrus (-50/-45/20) and additionally in the middle temporal gyrus, the inferior temporal gyrus, the supramarginal gyrus, the inferior parietal lobule, the parahippocampal gyrus, the lingual gyrus and the fusiform gyrus, **(C)** in the lower beta band (orange letter) in the bilateral cunei (-15/-80/25) and additionally in the precuneus, and **(D)** in the upper beta frequency band (red letter) in the bilateral precunei (0/-60/35) and slightly in the adjacent cingulate gyrus.

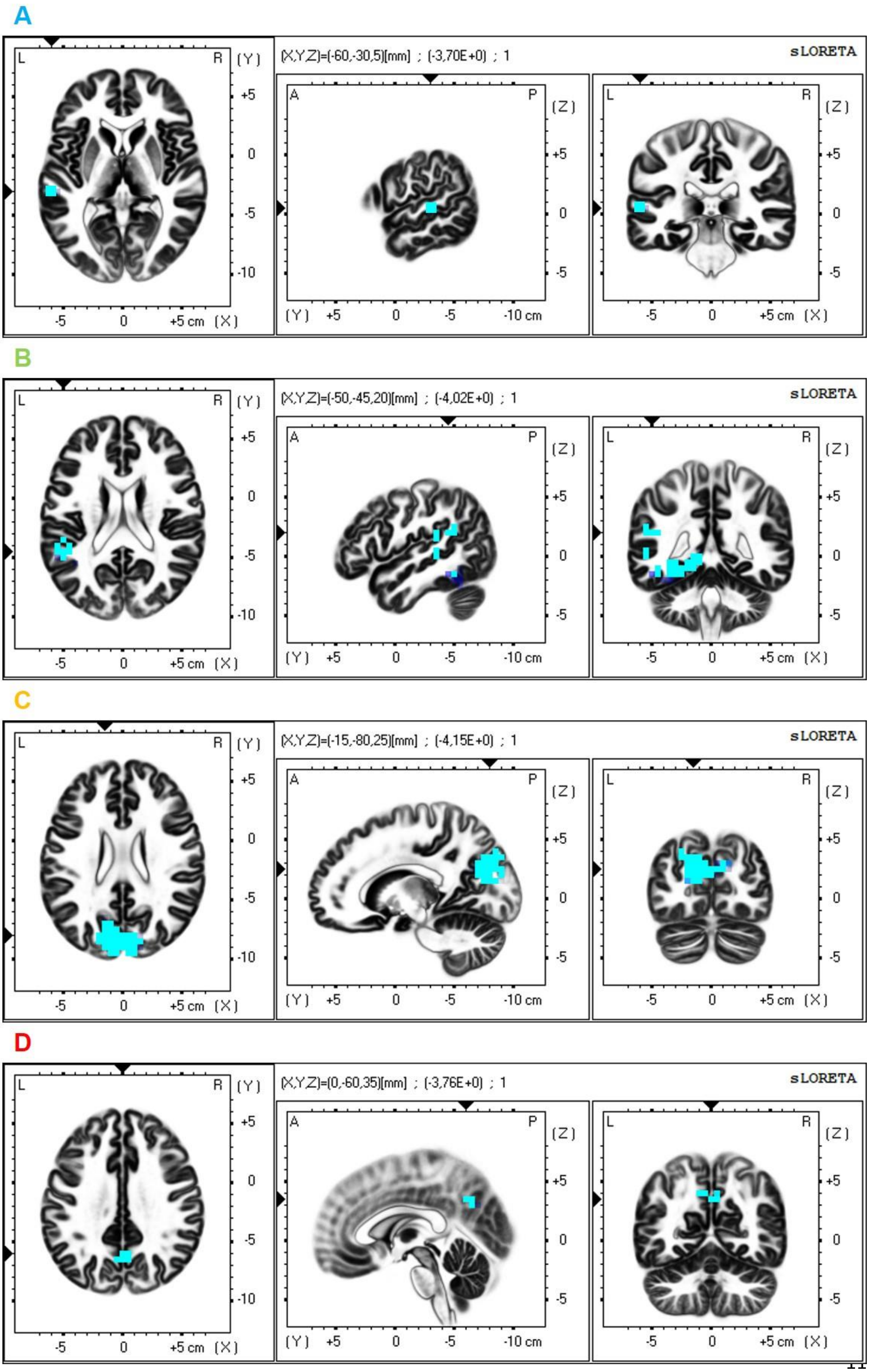
A
(X,Y,Z)=(-60,-30,5)[mm] ; (-3,70E+0) ; 1
sLORETA
B
(X,Y,Z)=(-50,-45,20)[mm] ; (-4,02E+0) ; 1
sLORETA
C
(X,Y,Z)=(-15,-80,25)[mm] ; (-4,15E+0) ; 1
sLORETA
D
(X,Y,Z)=(0,-60,35)[mm] ; (-3,76E+0) ; 1
sLORETA
L
R
A
P
(Y)
(Z)
(X)
-5
0
+5 cm
-10 cm

**Figure S6.** *Source-level amplitude effect based on to the a priori ROI-based analyses for the contrast signal versus noise trials*. The exact voxels of the *a priori* ROIs are marked in a turquoise color in the figure and their MNI coordinates are indicated in brackets in the text. In the time segment (3) before the response, power was significantly ($p < 0.05$) lower in signal than in noise trials in the lower beta frequency band (orange letters) in the **(A)** left cuneus (-10/-86/28) and **(B)** the left calcarine gyrus (-2/-82/20) as well as **(C)** in the upper beta frequency band (red letter) in the left precuneus (-8/-50/52).

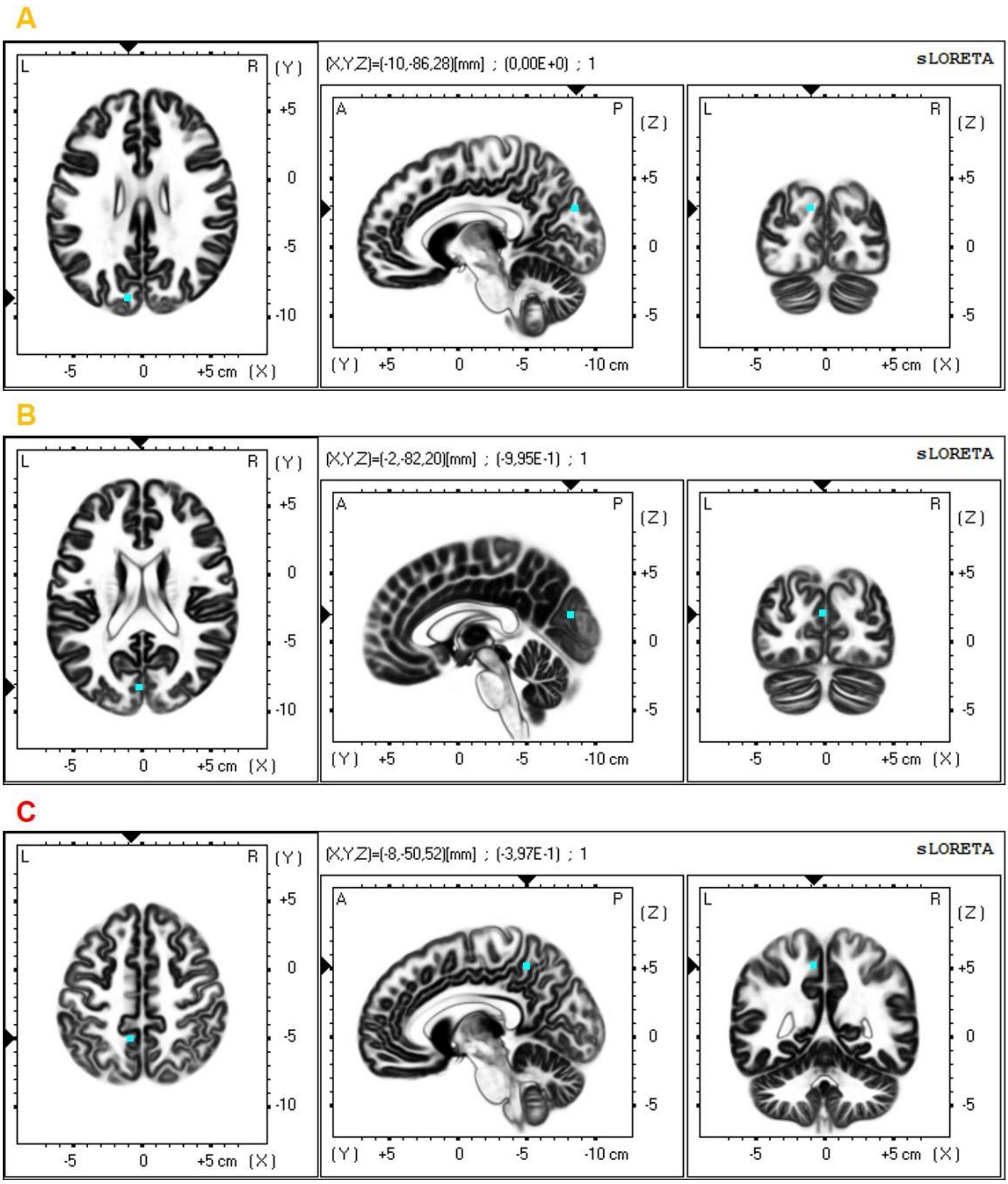
A
(X,Y,Z)=(-10,-86,28)[mm] ; (0,00E+0) ; 1
sLORETA
B
(X,Y,Z)=(-2,-82,20)[mm] ; (-9,95E-1) ; 1
sLORETA
C
(X,Y,Z)=(-8,-50,52)[mm] ; (-3,97E-1) ; 1
sLORETA

**Figure S7:** *Scalp-level amplitude effect for the contrast signal versus noise trials.*

**(A)** Theta band (blue letter): The coordinates -60/-30/5 of the left superior temporal gyrus (Figure S5A) correspond best to the scalp-level electrode T7 (see Table 1 in Scrivener and Reader (2022)). We calculated the same contrast with the scalp-level electrode T7 (indicated with a blue circle), which reached significance in the source-space for the left superior temporal gyrus (-60/-30/5) in the theta frequency band. In the time segment (3) before the response, power was significantly lower in signal than in noise trials in the theta frequency band at T7 ($p < 0.05$). The result is indicated with the turquoise filling of electrode T7 and is in line with the significant source-level effect illustrated in Figure S5A.

**(B)** Alpha band (green letter): The coordinates -50/-45/20 of the left superior temporal gyrus (Figure S5B) correspond best to the scalp-level electrode TP7 (see Table 1 in Scrivener and Reader (2022)). We calculated the same contrast with the scalp-level electrode TP7 (indicated with a green circle), which reached significance in the source-space in the alpha frequency band. In the time segment (3) before the response, power was significantly lower in signal than in noise trials in the alpha frequency band at TP7 ($p < 0.05$). The result is indicated with the turquoise filling of electrode TP7 and is in line with the significant source-level effect illustrated in Figure S5B.

**(C)** Lower beta band (orange letters): The coordinates -15/-80/25 (Figure S5C) and -10/-86/28 (Figure S6A) of the precuneus as well as -2/-82/20 of the left calcarine gyrus (Figure S6B) correspond best to the scalp-level electrode PO3 and $PO_Z$ (see Table 1 in Scrivener and Reader (2022)). We calculated the same contrast with the scalp-level electrodes PO3 and $PO_Z$ (indicated with an orange circle), which reached significance in the source-space in the lower beta frequency band. In the time segment (3) before the response, power was significantly lower in signal than in noise trials in the lower beta frequency band at PO3 and $PO_Z$ ($p < 0.05$). The result is indicated with the turquoise filling of electrodes PO3 and $PO_Z$ and is in line with the significant source-level effect illustrated in Figure S5C, Figure S6A and Figure S6B.

**(D)** Upper beta band (red letter): The coordinates 0/-60/35 (Figure S5D) and -8/-50/52 (Figure S6C) of the precuneus correspond to the scalp-level electrode $P_Z$ (see Table 1 in Scrivener and Reader (2022)). We calculated the same contrast with the scalp-level electrode $P_Z$ (indicated with a red circle), which reached significance in the source-space in the upper beta frequency band. In the time segment (3) before the response, power was significantly lower in signal than in noise trials in the upper beta frequency band at $P_Z$ ($p < 0.05$). The result is indicated with the turquoise filling of electrode $P_Z$ and is in line with the significant source-level effect illustrated in Figure S5D and Figure S6C.

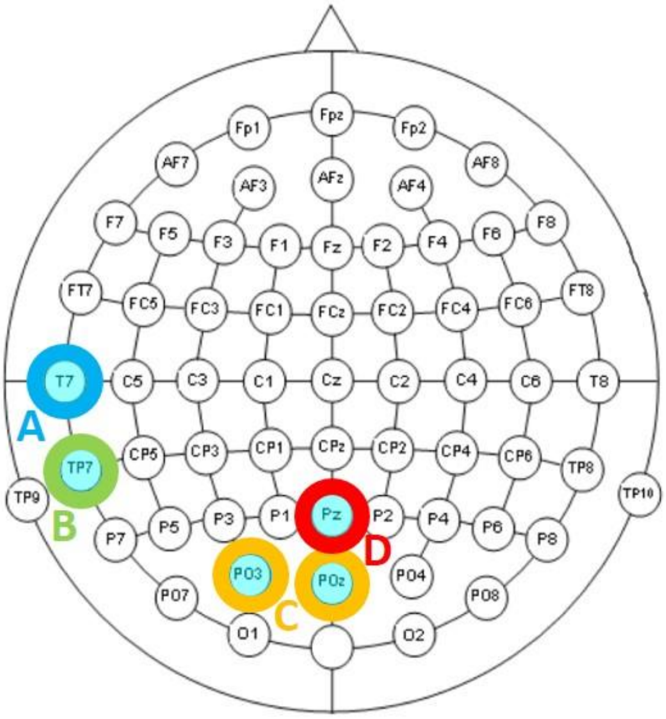
